# Extracellular vesicles and their miRNA cargo in retinal health and degeneration: mediators of homeostasis, and vehicles for targeted gene therapy

**DOI:** 10.1101/2020.03.29.014910

**Authors:** Yvette Wooff, Adrian V. Cioanca, Joshua A. Chu-Tan, Riemke Aggio-Bruce, Ulrike Schumann, Riccardo Natoli

## Abstract

1.1.

**Purpose:** Photoreceptor cell death and inflammation are known to occur progressively in retinal degenerative diseases, however the molecular mechanisms underlying these biological processes are largely unknown. Extracellular vesicles (EV) are essential mediators of cell-to-cell communication with emerging roles in the modulation of immune responses. EVs including exosomes encapsulate and transfer nucleic acids, including microRNA (miRNA), to recipient cells which in disease may result in dysfunctional immune responses and a loss of homeostatic regulation. In this work we investigated the role of isolated retinal small-medium sized EV (s-mEV) which includes exosomes in both the healthy and degenerating retina.

**Methods:** Isolated s-mEV from normal retinas were characterized using dynamic light scattering, transmission electron microscopy and western blotting, and quantified across 5 days of photo-oxidative damage-induced degeneration using nanotracking analysis. Small RNAseq was used to characterize the miRNA cargo of retinal s-mEV isolated from healthy and damaged retinas. Finally, the effect of exosome inhibition on cell-to-cell miRNA transfer and immune modulation was conducted using systemic daily administration of exosome inhibitor GW4869 and *in situ* hybridization of s-mEV-abundant miRNA, miR-124-3p. Electroretinography and immunohistochemistry was performed to assess functional and morphological changes to the retina as a result of GW4869-induced exosome depletion.

**Results:** Results demonstrated an inverse correlation between s-mEV secretion and photoreceptor survivability, with a decrease in s-mEV numbers following degeneration. Small RNAseq revealed that s-mEVs contained uniquely enriched miRNAs in comparison to in whole retinal tissue however, there was no differential change in the s-mEV miRNAnome following photo-oxidative damage. Exosome inhibition via the use of GW4869 was also found to exacerbate retinal degeneration, with reduced retinal function and increased levels of inflammation and cell death demonstrated following photo-oxidative damage in exosome-inhibited mice. Further, GW4869-treated mice displayed impaired translocation of photoreceptor-derived miR-124-3p to the inner retina during damage.

**Conclusions:** Taken together, we propose that retinal s-mEV and their miRNA cargo play an essential role in maintaining retinal homeostasis through immune-modulation, and have the potential to be used in targeted gene therapy for retinal degenerative diseases.

## 1.1. Introduction

Retinal degenerative diseases comprise a heterogeneous group of visual disorders associated with neuroinflammation and the progressive death of retinal neurons, often resulting in irreversible blindness^409^. Despite advances in understanding the pathogenesis and progression of retinal degenerative diseases ^3,18,19^, the precise molecular mechanisms that propagate retinal inflammation and subsequent cell death remain unknown. The study of extracellular vesicles (EV), including exosomes, might provide greater clarity in unpicking these mechanisms given their role as endogenous modulators of biological processes, including in inflammation ^408^, and intracellular communication pathways ^21,214,215^.

EV are small membrane-enclosed delivery vehicles which have been widely investigated for their vital role in mediating cell-to-cell communication in both healthy as well as diseased states^21,214,215^. Exosomes, the smallest EV fraction (40-200nm in diameter) ^207,211,407,410^, in particular, have recently been implicated in the pathogenesis of retinal degenerative diseases. These include Age-Related Macular Degeneration (AMD)^222,411-413^, Diabetic Retinopathy ^414^, and Retinitis Pigmentosa ^415^.

Exosomes are released by nearly all cell types and are formed via the endocytic pathway, with the invagination of the endosomal membrane allowing intraluminal vesicle formation in multivesicular bodies (MVB) ^213^. Biogenesis occurs either in an endosomal sorting complex required for transport (ESCRT)-dependent or ESCRT-independent manner ^222,223^, the latter of which can be blocked using GW4869, a non-competitive inhibitor of neutral sphingomyelinase 2 (nSMase2) – the key enzyme for generating exosomes via the ESCRT-independent pathway^226^. Following biogenesis, the MVB fuses with the host cell plasma membrane and releases the internalized exosomes into the extracellular environment ^204,207,213^. From here, exosomes can travel to nearby or distant target cells, often through biological fluids such as blood ^416^, to exert their biological effects ^204,205,417^. As exosomes selectively incorporate proteins, mRNA and non-coding RNA such as microRNA (miRNA) from their host cell (reviewed in ^204,205^), the transfer of exosomal contents can alter the environment of target recipient cells. In a healthy state, the transfer of exosomal nucleic acid contents, including miRNA, is required for homeostatic maintenance ^202,203^. However, in disease, aberrations to this process or the selective encapsulation of toxic proteins and/or dysregulated miRNA, can cause progressive inflammation ^236,237^.

MiRNA have been labelled as ‘master regulators’ of gene expression, due to their ability to target and repress multiple genes within and across different biological pathways ^197^. This regulatory power makes them ideal therapeutic and diagnostic molecules ^195^, in particular to combat dysregulated immune responses, such as those occurring in retinal degenerations ^196,197^. MiRNA have been reported to be enriched in exosomes over their host cells, suggesting that they are selectively incorporated to serve a dynamic, fast-response biological need ^200,201^.

The biological importance of exosomes and exosomal-miRNA (exoMiR) is reflected by their association with a range of inflammatory^132,418^, autoimmune^235,419^ and neurodegenerative diseases ^382,383,420^. These include diabetes ^421^, rheumatoid arthritis ^422-424^, and Alzheimer’s and Parkinson’s disease^236,420,425,426^. ExoMiR have been reported to play pathogenic roles in these diseases by promoting angiogenesis ^427,428^, and modulating immune responses, including in the recruitment of immune cells ^410,418,429-432^; features that contribute to cell death. To date however, the identification and role of exosomes and their miRNA cargo in the retina in both healthy and diseased states, is largely unexplored ^222,413^. While small extracellular vesicle fractions isolated following high speed >100,000xg ultracentrifugation and expressing tetraspanin markers CD9, CD63 and CD81 (such as those isolated in this work) are commonly referred to as exosomes, without evidence of endosomal origin and in complying with MISEV 2018 guidelines ^238^, are herein referred to as small-to-medium EV, or s-mEV. In reference to other works, EV terminology will be referred to as published in the original papers.

Characterizing the role of s-mEV and their miRNA cargo in both the normal and degenerating retina will aid in elucidating novel cell-to-cell communication pathways that could play a role in propagating inflammation during retinal degenerative diseases. Furthermore, uncovering the miRNA signature within retinal s-mEV as well as their potential binding partners may reveal novel regulatory mechanisms underpinning retinal degenerations, ultimately leading to the discovery of therapeutic targets. This study characterizes for the first time, retinal-derived s-mEV from both the healthy and degenerating mouse retina using a previously established model of photo-oxidative damage-induced retinal degeneration ^265^. Photo-oxidative damage models such as the one employed in this study accurately replicate key pathological changes seen in AMD, including the upregulation of oxidative stress and inflammatory pathways, progressive centralized focal photoreceptor cell loss and microglial/macrophage recruitment and activation ^264,265,272,273^.

We show that s-mEV secretion is inversely correlated to photoreceptor survivability, with the severity of retinal degeneration directly correlating to decreased retinal s-mEV numbers. We used small RNAseq to characterize the miRNA cargo of retinal s-mEV (exoMiR). Although we demonstrated that there was no change in s-mEV miRNA-cargo in response to retinal degeneration, we found that s-mEVs contain a set of uniquely enriched miRNAs. Further, we show that miRNA contained in retinal s-mEV were associated with the regulation of inflammatory, cell survival and motility pathways. Upon systemic exosome inhibition using GW4869, retinal function in healthy and photo-oxidative damaged mice was significantly reduced compared to controls. In addition, photoreceptor cell death and inflammation were significantly increased in GW4869-injected photo-oxidative damaged mice, compared to controls. Using *in situ* hybridization, we further demonstrated that the expression of miR-124-3p in the inner retina was reduced in GW4869-injected photo-oxidative damaged mice, suggesting that miR-124-3p movement could be mediated through s-mEV-dependent transport.

Taken together, these results suggest a novel role for s-mEV and s-mEV-miRNA-mediated cell-to-cell communication in the retina. We demonstrate that maintaining and transporting necessary levels of s-mEV cargo is required for normal retinal homeostasis and immunomodulation, with insufficient bioavailability of s-mEV potentially leading to inflammatory cell death (see Figure Error*! No text of specified style in document.*.*10*). In addition, this work elucidates downstream biological targets of s-mEV-miRNA that are required for retinal homeostatic maintenance, and identifies potential miRNA and mRNA therapeutic targets for further investigations.

## 1.2. Materials and Methods

### 1.2.1. Animal handling and photo-oxidative damage

All experiments were conducted in accordance with the ARVO Statement for the Use of Animals in Ophthalmic and Vision Research and with approval from the Australian National University’s (ANU) Animal Experimentation Ethics Committee (AEEC) (Ethics ID: A2017/41; Rodent models and treatments for retinal degenerations). Adult male and female C57BL/6J wild-type (WT) mice (aged between 50-90 postnatal days) were bred and reared under 12 h light/dark cycle conditions (5 lux) with free access to food and water. The C57BL/6J colony was genotyped for the presence of both the Rpe65^450Met^ polymorphism or the deleterious Crb1^rd8^ mutation using previously published primer sets ^275,276^. Sequencing for these was conducted at the ACRF Biomolecular Resource Facility, ANU. All animals used possessed the Rpe65^450Met^ polymorphism but were free of the Crb1^rd8^ mutation. Littermate age-matched WT mice were randomly assigned to photo-oxidative damage (PD) and dim-reared control (DR) groups. Animals in the photo-oxidative damage group were continuously exposed to 100k lux white LED light for a period of 1, 3, or 5 days as described previously ^265^, with the majority of experiments conducted for 5 days. Dim-reared control mice were maintained in 12h light (5 lux)/dark cycle conditions.

### 1.2.2. Retinal s-mEV isolation

Mice were euthanized with CO_2_ following experimental runs. Either two (from one mouse) or four (from two mice - used for high-throughput sequencing) retinas were pooled and collected in Hanks Buffered Saline Solution (HBSS, Gibco; Thermo Fisher Scientific, MA, USA). Retinas were transferred to 500μL digestion solution ((HBSS containing 2.5mg/mL papain (Worthington Biochemical, NJ, USA), 200U DNase I (Roche Diagnostics, NSW, AUS), 5μg/mL catalase (Sigma-Aldrich, MO, USA), 10μg/mL gentamycin (Sigma-Aldrich, MO, USA) and 5μg/mL superoxide dismutase (Worthington Biochemical, NJ, USA)) and finely chopped using scissors. Retinas were incubated at 37°C for 8 minutes, followed by 20 minutes at 8°C, to allow for the breakdown of the extracellular matrix and s-mEV release. Following digestion, tissue suspensions were neutralized by diluting in 11.5mL of HBSS and centrifuged at 1000xg for 10 minutes at 4°C to remove cells and cell debris. The supernatant was transferred to 14×89mm Beckman Ultra-Clear ultracentrifuge tubes (Beckman Coulter, CA USA) and centrifuged at 10,000xg for 30 minutes at 4°C in a Beckman Coulter Optima XE-100 (fitted with a SW41Ti Rotor (Beckman Coulter, CA USA)), to collect large EVs and remaining cell debris. The s-mEV-containing supernatant was transferred to new ultracentrifuge tubes and centrifuged for 1.5 hours at 150,000xg at 4°C. The supernatant was carefully decanted, and the s-mEV pellet resuspended via tituration for 1 minute in 500μL Ultrapure Endotoxin-free 0.1M PBS (Thermo Fisher Scientific, MA, USA) and used immediately for quantification.

For RNA isolation, the s-mEV pellet was resuspended immediately in 100μl RNAse A (10μg/ml in Ultrapure Endotoxin-free 0.1M PBS) and incubated for 30 minutes at 37°C to digest any RNA contamination. Following RNase treatment, s-mEV RNA was extracted using the mirVana miRNA Isolation Kit (Thermo Fisher Scientific, MA, USA) as per section 2.8.

### 1.2.3. s-mEV Characterization

#### 1.2.3.1. NanoSight

The size and concentration of s-mEV were measured using nanoparticle tracking analysis on a NanoSight NS300 (Malvern Instruments, Malvern, UK). s-mEV samples were diluted 1:20 (retinal s-mEV) or 1:40 (cell culture s-mEV) in 1ml Ultrapure Endotoxin-free 0.1M PBS to achieve a particle per frame value between 20 and 100. Samples were analyzed under constant flow provided by a syringe pump set at a speed of 35 (equating to ~3.1μl/min; ^433^. A total of nine 30s long videos were captured (camera setting: 14) for each sample. The detection threshold was set between 4-5 and was not altered between measurements of the same experiment. The concentration values, modal and mean sizes were exported to Prism V7 (GraphPad Software, CA, USA) for statistical analysis and plotting.

#### 1.2.3.2. Zetasizer

Dynamic light scattering measurements were performed using a Zetasizer Nano ZS 90 (Malvern Instruments, Malvern, UK). A 500μL undiluted retinal s-mEV suspension in Ultrapure Endotoxin-free 0.1M PBS was prepared, loaded in a low-volume disposable sizing cuvette (ZEN0112, Malvern Instruments, Malvern, UK) and agitated before measurements. Measurement parameters were set as follows: Material Refractive Index – 1.46, Dispersant Refractive Index – 1.330, Viscosity (cP) – 0.888, Temperature (°C) – 25, and Measurement Duration (s) – 60. The acquired intensity data was transformed using the General-Purpose Model within the Zetasizer analysis software to generate the size distribution of the s-mEV.

#### 1.2.3.3. Transmission Electron Microscopy (TEM)

A 30μl retinal s-mEV suspension was placed on a 200-mesh carbon-coated copper grid (Sigma-Aldrich, MO USA) and pre-treated with glow discharge using an Emtech K100X system (Quorum Technologies, Sussex, UK). After 20 minutes, s-mEV were contrasted with 2% uranyl acetate solution for 1 minute, followed by 3 washes in 0.22μm filtered PBS (Thermo Fisher Scientific, MA, USA). Excess PBS was removed by placing a piece of absorbent paper at the edge of the grid. The grids were imaged on a Hitachi 7100 FA transmission electron microscope (Hitachi, Tokyo, Japan) at 100kV. The images were captured with a side mounted Gatan Orius CCD camera (Gatan, CA, USA) at 4008×2672 pixels resolution using a 2 seconds exposure operated through Gatan Microscopy Suite (Gatan, CA, USA). A total of 20 images were captured at 100,000x magnification from four different grids, each containing s-mEV isolated from a different retinal s-mEV preparation (2 mouse retinas/preparation). The images were imported into ImageJ V2.0 software (National Institutes of Health, Bethesda, MD, USA), scale-calibrated and the diameter of approximately 230 s-mEV was measured. The size distribution was plotted in a histogram with 20nm wide bins using Prism V7.0 (GraphPad Software, CA, USA).

#### 1.2.3.4. Western blot

s-mEV pellets (see section 2.2) were immediately lysed in 50μL CellLyticTM Cell Lysis Buffer (supplemented with 1:100 protease inhibitor cocktail; Sigma-Aldrich, MO, USA). The blot was performed as previously described ^67^. Briefly,10μg of denatured protein was loaded onto Novex 4-20% Tris-Glycine Mini Gels (Thermo Fisher Scientific, MA, USA) and subjected to electrophoresis (45 minutes, 150V). The protein bands were transferred (45 minutes, 20V) to a nitrocellulose membrane (Bio-Rad, CA, USA) using the Power Blotter semi-dry system (Thermo Fisher Scientific, MA, USA). Membranes were then washed in PBS-Tween (0.025%; PBS-T), blocked in 3% BSA for 1 hour and then incubated overnight at 4°C with primary s-mEV marker antibodies CD63, (1:1000, Ts63, Thermo Fisher Scientific, MA, USA), CD81 (1:2000, ab109201, Abcam, Cambridge, UK) or CD9 (1:2000, ab92726, Abcam, Cambridge, UK). Following three washes in PBS-T, blots were incubated in appropriate secondary antibodies, HRP-conjugated Goat Anti-Rabbit IgG (H+L) (1:1000, 170-6515, Bio-Rad, CA, USA) or Goat-anti-Mouse IgG (1:1000, 170-6516, Bio-Rad, CA, USA) for 2 hours at room temperature. Membranes were washed in PBS-T and developed for 2 minutes with ClarityTM Western ECL Substrate (Bio-Rad, CA, USA). Imaging was performed using a ChemiDocTM MP Imaging System with Image Lab^TM^ software (Bio-Rad, CA, USA).

### 1.2.4. Exosome inhibition

Exosome inhibition was performed using GW4869 (Sigma-Aldrich, MO, USA), a known inhibitor of exosome biogenesis and release ^434^. GW4869 was reconstituted in dimethyl sulfoxide (DMSO; Sigma-Aldrich, MO, USA) to a concentration of 5mM and used as a stock solution for further dilution in Ultrapure Endotoxin-free 0.1M PBS. Mice were injected with 1.25mg/kg GW4869 via intraperitoneal (I.P.) injection daily for 5 days. 10.3% DMSO in Ultrapure Endotoxin-free 0.1M PBS (corresponding to the final volume of DMSO in GW4869 preparations) was used as a negative control. All mice were monitored daily for signs of distress or sickness.

### 1.2.5. Retinal Assessment

#### 1.2.5.1. Retinal Function via Electroretinography (ERG)

To assess retinal function full-field scotopic ERG was performed as previously described ^17^. Briefly, mice were dark-adapted overnight before being anaesthetized with an intraperitoneal injection of Ketamine (100 mg/kg; Troy Laboratories, NSW, Australia) and Xylazil (10 mg/kg; Troy Laboratories, NSW, Australia). Both pupils were dilated with one drop each of 2.5% w/v Phenylephrine hydrochloride and 1% w/v Tropicamide (Bausch and Lomb, NY, USA).

Anaesthetized and pupil dilated mice were placed on the thermally regulated stage of the Celeris ERG system (Diagnosys LLC, MA, USA). The Celeris ERG system has combined Ag/AgCl electrode-stimulator eye probes which measure the response from both eyes simultaneously, and uses 32-bit ultra-low noise amplifiers fitted with impedance testing. Eye probes were cleaned with 70% ethanol and then a 0.3% Hypromellose eye drop solution (GenTeal; Novartis, NSW, AUS) was applied to both probes. The probes were then placed covering and just touching the surface of each eye. A single- or twin-flash paradigm was used to elicit a mixed response from rods and cones. Flash stimuli for mixed responses were provided using 6500K white flash luminance range over stimulus intensities from −0.01 − 40 log cd s m^−2^. Responses were recorded and analyzed using Espion V6 Software (Diagnosys LLC, MA, USA). Statistics were performed in Prism V7.0 using a two-way analysis of variance (ANOVA) to test for differences in a-wave and b-wave responses. Data was expressed as the mean wave amplitude ± SEM (μV).

#### 1.2.5.2. Optical Coherence Tomography (OCT)

Cross-sectional images of live mouse retinas were taken at 1mm increments from the optic nerve using a Spectralis HRA+OCT device (Heidelberg Engineering, Heidelberg, Germany) as previously described ^265^. Eye gel (GenTeal; Novartis, NSW, AUS) was administered to both eyes for recovery.

Using OCT cross-sectional retinal images, and ImageJ V2.0 software (National Institutes of Health, Bethesda, MD, USA), the thickness of the outer nuclear layer (ONL), was calculated as the ratio to the whole retinal thickness (outer limiting membrane to the inner limiting membrane).

### 1.2.6. Retinal tissue collection and preparation

Animals were euthanized with CO_2_ following functional ERG analysis. The superior surface of the left eye was marked and enucleated, then immersed in 4% paraformaldehyde for 3 hours. Eyes were then cryopreserved in 15% sucrose solution overnight, embedded in OCT medium (Tissue Tek, Sakura, Japan) and cryosectioned at 12μm in a parasagittal plane (superior to inferior) using a CM 1850 Cryostat (Leica Biosystems, Germany). To ensure accurate comparisons were made for histological analysis, only sections containing the optic nerve head were used for analysis. The retina from the right eye was excised through a corneal incision and placed into RNAlater solution (Thermo Fisher Scientific, MA, USA) at 4°C overnight and then stored at −80°C until further use.

### 1.2.7. Immunolabelling

Immunohistochemical analysis of retinal cryosections was performed as previously described ^53^. Fluorescence was visualized and images taken using a laser-scanning A1^+^ confocal microscope at 20x magnification (Nikon, Tokyo, Japan). Images panels were analyzed using ImageJ V2.0 software and assembled using Photoshop CS6 software (Adobe Systems, CA, USA).

#### 1.2.7.1. IBA-1 Immunohistochemistry

Immunolabeling for IBA-1 (1:500, 019-19741, Wako, Osaka, Japan) and quantification was performed as previously described ^53^. The number of IBA-1^+^ cells (a marker of retinal microglia and macrophages) was counted across the superior and inferior retina using two retinal sections per mouse and then averaged. Retinal cryosections were stained with the DNA-specific dye bisbenzimide (1:10000, Sigma-Aldrich, MO, USA) to visualize the cellular layers.

#### 1.2.7.2. TUNEL assay

Terminal deoxynucleotidyl transferase (Tdt) dUTP nick end labelling (*TUNEL*), was used as a measure of photoreceptor cell death. TUNEL *in situ* labelling was performed on retinal cryosections using a Tdt enzyme (Cat# 3333566001, Sigma-Aldrich, MO, USA) and biotinylated deoxyuridine triphosphate (dUTP) (Cat# 11093070910, Sigma-Aldrich, MO, USA) as previously described ^286^. Images of TUNEL staining were captured with the A1^+^ Nikon confocal microscope at 20x magnification. The total number of TUNEL^+^ cells were counted including both the superior and inferior retina using two retinal sections per animal, and is represented as the average number of TUNEL^+^ cells per retinal section.

To further quantify photoreceptor survival, the thickness of the ONL on retinal cryosections was determined by counting the number of nuclei rows (photoreceptor cell bodies) in the area of retinal lesion development (1mm superior to the optic nerve head). Photoreceptor cell row quantification was performed five times per retina using two retinal cryosections at comparable locations per mouse. The thickness the ONL, inner nuclear layer (INL), and the combined ganglion cell layer (GCL)-outer plexiform layer (OPL) thickness were also measured at the lesion site on the superior retina, and expressed as a ratio to whole retinal thickness.

#### 1.2.7.3. In situ hybridization

Localization of miR-124-3p within the retina was determined by *in situ* hybridization. A double DIG-labelled miR-124-3p miRCURY LNA miRNA Detection Probe (Exiqon, Vedbaek, Denmark) was used on retinal cryosections, which were hybridized for 1 hour at 53°C as previously described ^245^. The bound probe was visualized using 5-bromo-4-chloro-3 indoyl phosphate (NBT/BCIP; Sigma-Aldrich Corp., St. Louis, MO, USA). Bright field images were captured on the A1^+^ Nikon confocal microscope fitted with a DS-Ri1-U3 color camera at 20x magnification and 4076×3116 pixel resolution. All images were centered at the site of lesion located approximately 1mm superiorly to the optic nerve head. The images were imported into ImageJ V2.0 software, converted to 8-bit format and then the densitometry was calculated. Mean grey values were measured at five different locations along the INL, ONL and the outer limiting membrane/photoreceptor inner segment region with background levels subtracted prior.

### 1.2.8. RNA extraction

RNA (enriched for miRNA) extraction and purification from retinas or RNase A treated s-mEV pellets was performed using an acid-phenol:chloroform extraction method with the mirVana miRNA Isolation Kit (Thermo Fisher Scientific, MA, USA) according to the manufacturer's instructions. The concentration and purity of each RNA sample was assessed using the ND-1000 spectrophotometer (Nanodrop Technologies, DE, USA). The size distribution and concentration of s-mEV miRNA was further assessed using a 2100 Agilent Bioanalyzer with an Agilent Small RNA Kit (Agilent Technologies, CA, USA), according to the manufacturers’ instruction.

#### 1.2.8.1. cDNA synthesis from mRNA and miRNA templates

Following purification of RNA, cDNA was synthesized from 1μg RNA using either the Tetro cDNA Synthesis Kit (Bioline Reagents, London, UK) from an mRNA template, or using the TaqMan MicroRNA RT kit (Thermo Fisher Scientific) from a miRNA template, according to manufacturers’ instructions.

#### 1.2.8.2. Quantitative real-time polymerase chain reaction

The expression of ESCRT-independent exosome biogenesis pathways genes was measured by qRT-PCR. We targeted *Pdcd6ip* (also known as *Alix*), which encodes an accessory protein in the ESCRT-dependent pathway, and *Smpd3*, which encodes nSMase2 in the ESCRT-independent pathway ^207^. The expression of miR-124-3p was also investigated in retinal lysates from exosome-inhibited mice, and controls. The expression of these genes and miRNA was measured using mouse specific TaqMan hydrolysis probes (Table Error! No text of specified style in document..*1*) and TaqMan Gene Expression Master Mix (Thermo Fisher Scientific, MA, USA). Reactions were performed in technical duplicates in a 384-well format using a QuantStudio 12 K Flex RT-PCR machine (Thermo Fisher Scientific, MA, USA). Data was analyzed using the comparative C_t_ method (ΔΔC_t_) and results are presented as percent change relative to control. Expression was normalized to reference gene glyceraldehyde-3-phosphate dehydrogenase (*Gapdh*) for mRNA, and small nuclear RNA U6 for miRNA.

**Figure Error! No text of specified style in document..1:**
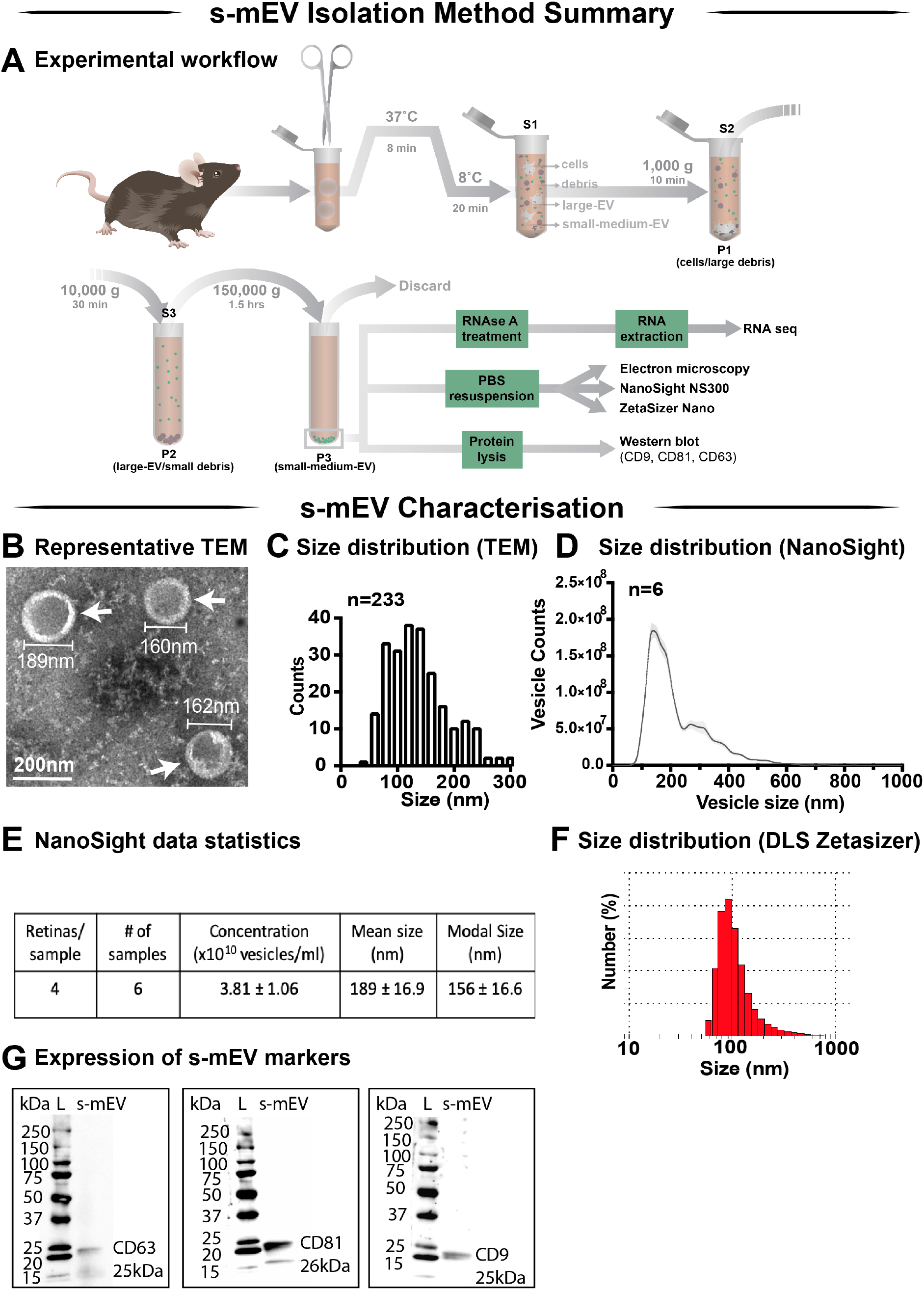
Retinal s-mEV isolation and characterization. **(A)** Workflow for the isolation and characterization of retinal s-mEV. s-mEV were isolated from papain-digested retinas using differential ultracentrifugation. s-mEV pellets (P3) were processed for RNA extraction and RNA sequencing (RNA seq) and size and morphology analysis by electron microscopy, NanoSight NS300 and Zetasizer Nano. s-mEV pellets were also digested in protein lysis buffer and analyzed for the expression of s-mEV markers CD9, CD81 and CD63 by western blot. S=supernatant, P=Pellet. **(B)** Representative Transmission Electron Microscopy (TEM) micrograph showing isolated retinal vesicles (white arrows) with consistent round, cup-shaped morphology and size of s-mEV. **(C)** Diameter distribution of retinal s-mEV measured from TEM micrographs (n=233). **(D)** Size distribution of retinal s-mEV (n=6 samples, 4 retinas/sample) using nanotracking analysis performed on a NanoSight NS300. **(E)** Average concentration, mean and modal size of s-mEV isolated from mouse retinas measured by nanotracking analysis (n=6 samples, 4 retinas/sample). **(F)** Representative plot of the size distribution of s-mEV measured by dynamic light scatter (DLS) using a Zetasizer Nano instrument (n=5). **(G)** Full-length western blots showing the presence of s-mEV markers CD63 (25kDa), CD9 (26kDa) and CD81 (25kDa) in retinal s-mEV protein lysates (Exo). Molecular weight standard (L). Scale bar = 200nm.

**Table Error! No text of specified style in document..1:** TaqMan Hydrolysis probes (Thermo Fisher Scientific, MA, USA) used for qRT-PCR.

| Gene symbols | Gene name | Catalogue number |
| --- | --- | --- |
| *miR-124a* | Mmu_miR-124-3p | 001182 (assay ID) |
| *U6 snRNA* | Small nuclear RNA U6 | 001973 (assay ID) |
| *Gapdh* | Glyceraldehyde-3-Phosphatase Dehydrogenase | Mm01536933_m1 |
| *Pdcd6ip* | Programmed Cell Death 6 Interacting Domain | Mm00478032_m1 |
| *Smpd3* | Sphingomyelin Phosphodiesterase 3 | Mm00491359_m1 |

### 1.2.9. Small RNA high-throughput sequencing

#### 1.2.9.1. Library preparation and sequencing

cDNA libraries from miRNA-enriched s-mEV RNA samples were prepared by the John Curtin School of Medical Research Biomolecular Research Facility (JCSMR, BRF, ACT, AUS), using the Capture and Amplification by Tailing and Switching method (CATS RNA-seq Kit v2 x24, Diagenode Cat# C05010041, Leige, Belgium). 10ng RNA was used as input and the dephosphorylation step omitted to select for 3’-OH RNA species (miRNA). The library was amplified with 9 PCR cycles and cleaned with 0.9x AMPure^®^ XP beads (A63881, Beckman Coulter, CA, USA) to enrich for DNA fragments shorter than 50nt. Libraries were multiplexed and sequenced on a single lane using the Illumina NextSeq500 (Illumina, CA, USA) acquiring 50 base-pairs single-end reads. The sequencing depth was between 8 and 15 million reads/sample with an average phred read quality of 33 (Figure S3 A). Sequencing libraries prepared from whole retinal tissue were retrieved from BioProject database (PRJNA606092). These libraries were previously prepared by our group using the same library construction method, and the same bioinformatic analysis pipeline was applied as stated below, see Section 2.9.2.

#### 1.2.9.2. Bioinformatics

Sequencing reads were initially checked for quality scores, adapter/index content and K-mer content using FastQC v0.11.8 (Babraham Bioinformatics, Cambridge, UK), then imported into Partek^®^ Flow^®^ (Partek Inc, MO, USA) for all subsequent analyses. Base composition analysis indicated the presence of an enriched poly-A tail and the CATS template-switching nucleotide within the first 3 base-pairs of the reads (Figure S3 B), indicating a successful library preparation. A trimming pipeline was created according to the following cutadapt code “cutadapt −-trim-n −a GATCGGAAGAGCACACGTCTG −a AGAGCACACGTCTG <input.file> | cutadapt −u 3 −a AAAAAAAAAACAAAAAAAAAA −e 0.16666666666666666 −| cutadapt −g GTTCAGAGTTCTACAGTCCGACGATC −m 18 −o <output.file> −”, to remove the template switching nucleotide and the 3’ and 5’ adapters associated with the CATS library preparation. After trimming, the base composition was centered around 25% for each nucleotide (Figure S3 C), and FastQC analysis confirmed the effective removal of the 3’ and 5’ adapters and indices.

Trimmed reads were aligned against the mature mouse miRNA downloaded from miRbase v.22 using the Burrows-Wheeler Aligner (BWA) in backtrack mode with parameters −n 1(base), −o 1, −e −1, −d 10, −i 5, −k 2, −E 4, −R 30, −t 24 (as recommended by ^296^. MiRNAs with less than 10 alignments across all samples were discarded from the subsequent analyses. Reads aligned to miRbase v.22 (mature miRNAs) with BWA had a length distribution between 18-24 nucleotides consistent with the expected length of mature miRNA (Figure S3 D). Aligned reads were normalized using the Trimmed means of M (TMM) and Upper Quartile (UQ) methods as recommended ^297^, with the latter being chosen as the preferred method as it produced less variable means and distributions (Figure S3, E, F and G). After normalization, a two-dimensional principal component analysis (PCA) was performed in Partek^®^ Flow^®^ (Partek Inc, MO, USA) to assess the clustering of the samples and identify outliers. Fold changes and statistical significance were computed using the Gene Specific Analysis (GSA) tool within Partek^®^ Flow^®^ (Partek Inc, MO, USA). This function uses the corrected Akaike Information Criterion to select the best statistical model for each gene from the available Normal, Negative Binomial, Lognormal or Lognormal with shrinkage options. Lognormal with shrinkage ^298^, produced the best fit and thus was selected to perform differential analysis. A dataset was created containing the expression values for each miRNA in all samples and imported into R ^299^. The counts were log2 transformed to display the distribution of miRNAs in dim-reared and photo-oxidative samples using the packages ggbeeswarm ^300^ and ggplot2 ^301^. Hierarchical clustering analysis (HCA) was performed in Partek^®^ Flow^®^ (Partek Inc, MO, USA) using the Euclidian distance as a point metric distance and the average linkage as the agglomerative method.

Sequencing data can be accessed from BioProject (Accession ID: PRJNA615966).

#### 1.2.9.3. Network and pathway enrichment analysis

The miRNet platform ^302^ was used to elucidate potential interactions between exosomal miRNA and retinal mRNA. The mRNA dataset is available from BioProject (accession ID: PRJNA606092) and comprises of all retinal genes with a log_2_(count per million) value > −2.4 (Table S2). The retinal targetome of the top 10 exosomal miRNAs as well as the targetome of the s-mEVs enriched miRNA were separately imported into Enrichr ^303^ and analyzed for over-expressed pathways annotated in Wikipathways ^304^ (mouse annotation), and gene-disease associations listed in DisGeNet database ^305^.

### 1.2.10. *In vitro* experiments

#### 1.2.10.1. 661W cell culture

Murine photoreceptor-derived 661W cells (kindly gifted by Dr. Muayyad R. Al-Ubaidi, Dept. of Biomedical Engineering, University of Houston, Houston, TX, USA) ^291^, were used for *in vitro* experiments at passage 1-5. The authenticity of the of the cells was validated by short tandem repeat analysis (CellBank, Sydney, AUS). Cells were cultured in growth media (Dulbecco’s Modified Eagle Medium (DMEM; Sigma-Aldrich, MO, USA) supplemented with 10% fetal bovine serum (FBS; Sigma-Aldrich, MO, USA), 6 mM L-glutamine (Thermo Fisher Scientific, MA, USA) and antibiotic-antimycotic (100U/ml penicillin, 100 μg/ml streptomycin; Thermo Fisher Scientific, MA, USA)), as previously published ^295^.

Cells were maintained and all incubation steps were performed in dark conditions in a humidified atmosphere of 5% CO_2_ at 37°C, unless otherwise stated. Cells were passaged by trypsinization every 3 – 4 days.

To deplete FBS of s-mEV, the serum was centrifuged (200,000xg, at 4^°^C for 18 hours) using a Beckman Coulter Optima XE-100 Ultracentrifuge (Beckman Coulter, CA, USA), with a SW41Ti rotor (Beckman Coulter, CA USA) and the supernatant used as FBS supplement in all GW4869 experiments (adjusted growth media).

#### 1.2.10.2. In vitro photo-oxidative damage

661W cells were seeded in 96 well plates (Nunc, Thermo Fisher Scientific, MA, USA) at 2×10^4^ cells/well 24 hours prior to photo-oxidative damage experiments. Cells were exposed for 4 hours to 15,000 lux light (2.2 mW/cm^2^; irradiance measured with PM100D optical power meter, THORLABS, NJ, USA) from two white fluorescent lamps (2 × 10W T4 tri-phosphor 6500K daylight fluorescent tubes; Crompton, NSW, Australia) as published previously ^322,435^. Control cells were completely wrapped in aluminum foil with six small incisions to allow air/gas exchange.

#### 1.2.10.3. Cell viability by MTT assay

Cell viability was tested by 3-(4,5-dimethylthiazol-2-yl)-2,5-diphenyltetrazolium bromide (MTT) assay (Roche, Mannheim, Germany), according to manufacturer’s instructions. Briefly, conditioned media from 661W cells undergoing photo-oxidative damage or grown under dim conditions and treated with GW4869 at 5, 10, 20 and 40μM concentrations or equivalent concentrations of DMSO was discarded and 100μl growth media containing 10% MTT was added to all wells. The cells were incubated for 4 hours to allow for the formation of formazan crystals then 100μl of solubilization buffer was added to each well and cells incubated for 18 hours. The absorbance at 570nm of each well was measured with a 670nm reference wavelength using an Infinite® 200 Pro plate reader (Tecan, Mannedorf, Switzerland). Background absorbance (wells containing medium with MTT reagent only) was subtracted from all sample wells and the viability calculated by dividing the absorbance of treated wells (DMSO or GW4869) by controls (cell treated with medium only).

#### 1.2.10.4. Exosome inhibition in 661W cells

To test the effect of GW4869 on exosome inhibition, 661W cells were seeded at a density of 1×10^5^ cells/well in 6 well plates (Nunc, Thermo Fisher Scientific, MA, USA) and grown to 80% confluence in growth medium. Media was removed and cells treated with 20μM GW4869, or equivalent DMSO, in adjusted growth media for 4h. The conditioned medium from four GW4869-treated or control wells (661W cells treated with equivalent DMSO concentration) was pooled and processed for s-mEV isolation and characterization as described in Sections 2.2 and 2.3.1.

#### 1.2.10.5. Statistical analyses

All graphing and statistical analyses were performed using Prism V7.0, unless otherwise specified. An unpaired Student’s *t*-test, one-way analysis of variance (ANOVA), or two-way ANOVA with Tukey’s multiple comparison post-hoc test was utilized as appropriate to determine the statistical outcome. *Non-adjusted p values (P<0.05) or false-discovery adjusted p values (FDR<0.1) were deemed statistically significant*. All data was expressed as the mean ± SEM.

## 1.3. Results

### 1.3.1. Retinal s-mEV isolation

In this work we demonstrate a novel protocol for the isolation of s-mEV from mouse retinas (Figure Error*! No text of specified style in document.*.***1***A). Extracellular vesicles isolated from dim-reared retinas displayed properties of s-mEV, including the distinctive round, cup-shaped morphology of exosomes ^436^ as seen in representative negative-stained TEM images (Figure Error*! No text of specified style in document.*.***1***B, and Supplementary Figure Error! No text of specified style in document.**.1**A). Isolated vesicles were within the expected size-range of s-mEV including exosomes ^207,211,238,410^, as shown by TEM size distribution histogram, nanotracking analysis using NanoSight NS300 and dynamic light scatter using a Zetasizer Nano instrument (Figure Error*! No text of specified style in document.*.***1***C-F). Further, isolated vesicles displayed known s-mEV markers CD63, CD81 and CD9, as determined by western blotting (Figure Error*! No text of specified style in document.*.***1***G).

### 1.3.2. Retinal s-mEV decrease in concentration but not size, during photo-oxidative damage-induced degeneration

Following the characterization of retinal s-mEV isolated from dim-reared retinas, the effect of photo-oxidative damage on the secretion of s-mEV was investigated (Figure Error*! No text of specified style in document.*.***2***A). The relative retinal s-mEV concentration was found to decrease in a damage-dependent manner, reducing significantly from dim-reared controls by 17±2.7% after up to 3 days of photo-oxidative damage and 38±4.0% after 5 days of photo-oxidative damage (Figure Error*! No text of specified style in document.*.***2***B). While the s-mEV concentration after 5 days of photo-oxidative damage was decreased from (4.27±0.2) ×10^10^ vesicles/ml to (2.82±-0.3) x10^10^ vesicles/ml (Figure Error*! No text of specified style in document.*.***2***C). The analysis of the size distribution of s-mEV at these two timepoints showed that vesicles with diameter of up to 200nm had a more pronounced decrease in numbers (Figure Error*! No text of specified style in document.*.***2***D). The mean and modal size of s-mEV remained unchanged after 5 days of photo-oxidative damage (Figure Error*! No text of specified style in document.*.***2***E). Taken together these results suggest that while there is no change in the size of isolated s-mEV, there was a progressive reduction in the concentration of exosomes during photo-oxidative damage-induced degeneration.

**Figure Error! No text of specified style in document..2:**
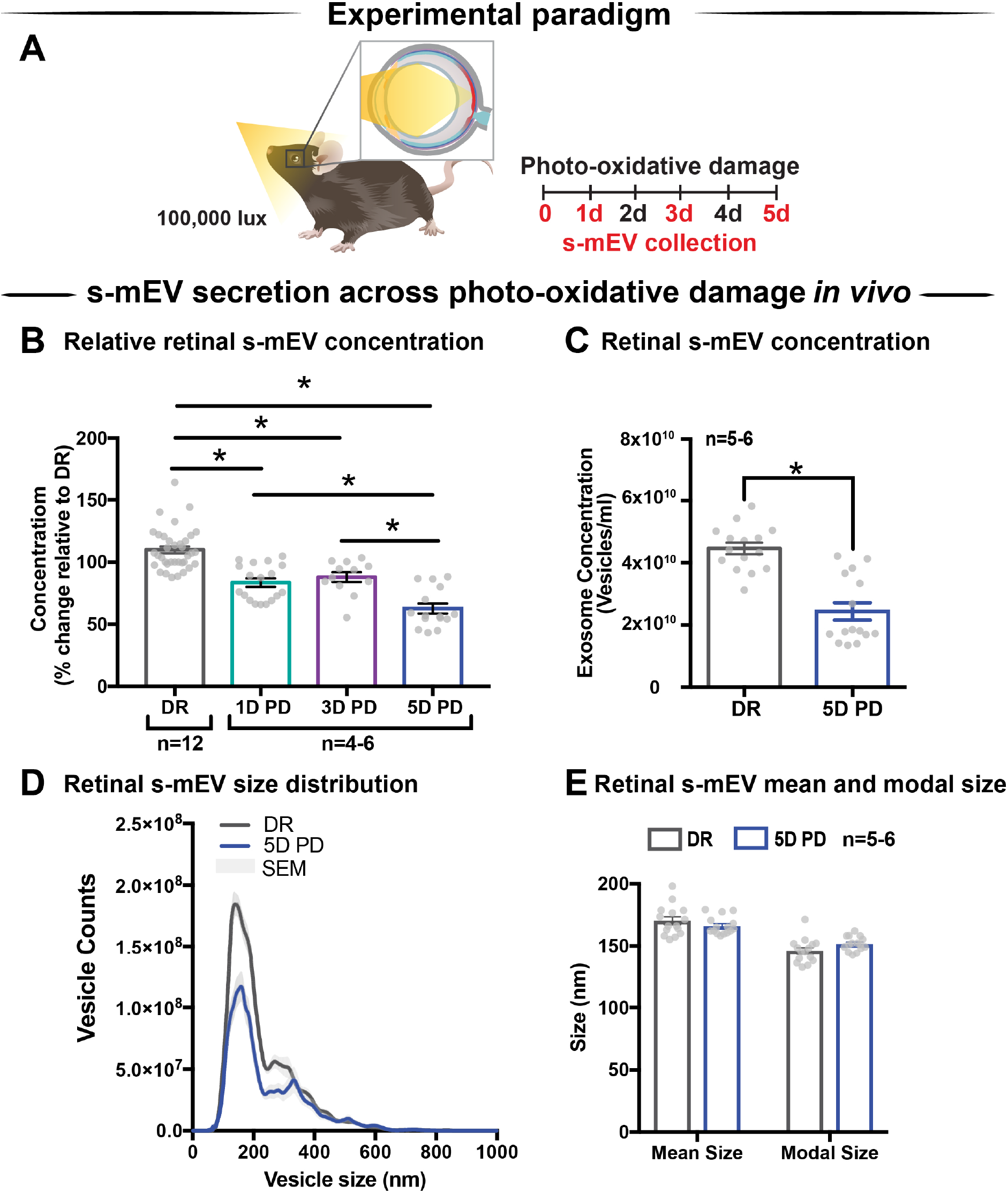
Retinal s-mEV decrease in concentration but not size, over photo-oxidative damage-induced degeneration. **(A)** Experimental paradigm, with red retinal labelling indicating lesion site. **(B)** Concentration of retinal s-mEV decreased progressively across time in PD, with significantly reduced s-mEV at 1D, 3D and 5D PD compared to DR controls (P<0.05, n=3-12). Concentration values are presented as percentage of DR. **(C)** Significant reduction in the concentration of s-mEV from 5D PD retinas compared to DR controls (P<0.05, n=5-6). **(D**) No significant change in the size distribution of s-mEV isolated from 5D PD retinas compared to DR controls, however vesicles with size < 200nm show a marked decrease in concentration following 5D PD (P>0.05, n=5-6). **(E)** No difference in either the mean or modal size of retinal s-mEV between DR and 5D PD groups (P>0.05, n=5-6). All results obtained using nanotracking analysis (NanoSight NS300).

### 1.3.3. GW4869 inhibits s-mEV bioavailability in vitro

Given the correlation between increased photoreceptor cell death and reduced retinal s-mEV concentration during photo-oxidative damage, the effect of s-mEV, specifically exosome inhibition was investigated using GW4869. To determine efficiency, GW4869 was added in culture to 661W photoreceptor-like cells (Figure Error*! No text of specified style in document.*.***3***A). s-mEV isolated from 661W were characterized using TEM (Figure Error*! No text of specified style in document.*.***3***B and Supplementary Figure Error! No text of specified style in document.**.1**B). Compared to DMSO controls, 661W s-mEV concentration (Figure Error*! No text of specified style in document.*.***3***C-D) was significantly reduced in GW4869-treated cells, demonstrating a high level of efficacy of this inhibitor. Furthermore, increased GW4869 concentrations did not affect cell viability compared to DMSO controls under dim conditions. However, following photo-oxidative damage (4 hours of 15,000k lux white light), an inverse dose-dependent correlation was observed between GW4869 concentration and cell viability (Figure Error*! No text of specified style in document.*.***3***E-F). Overall GW4869 was shown to reduce the bioavailability of isolated s-mEV *in vitro,* with the reduction likely attributed to a decrease in exosome production.

**Figure Error! No text of specified style in document..3:**
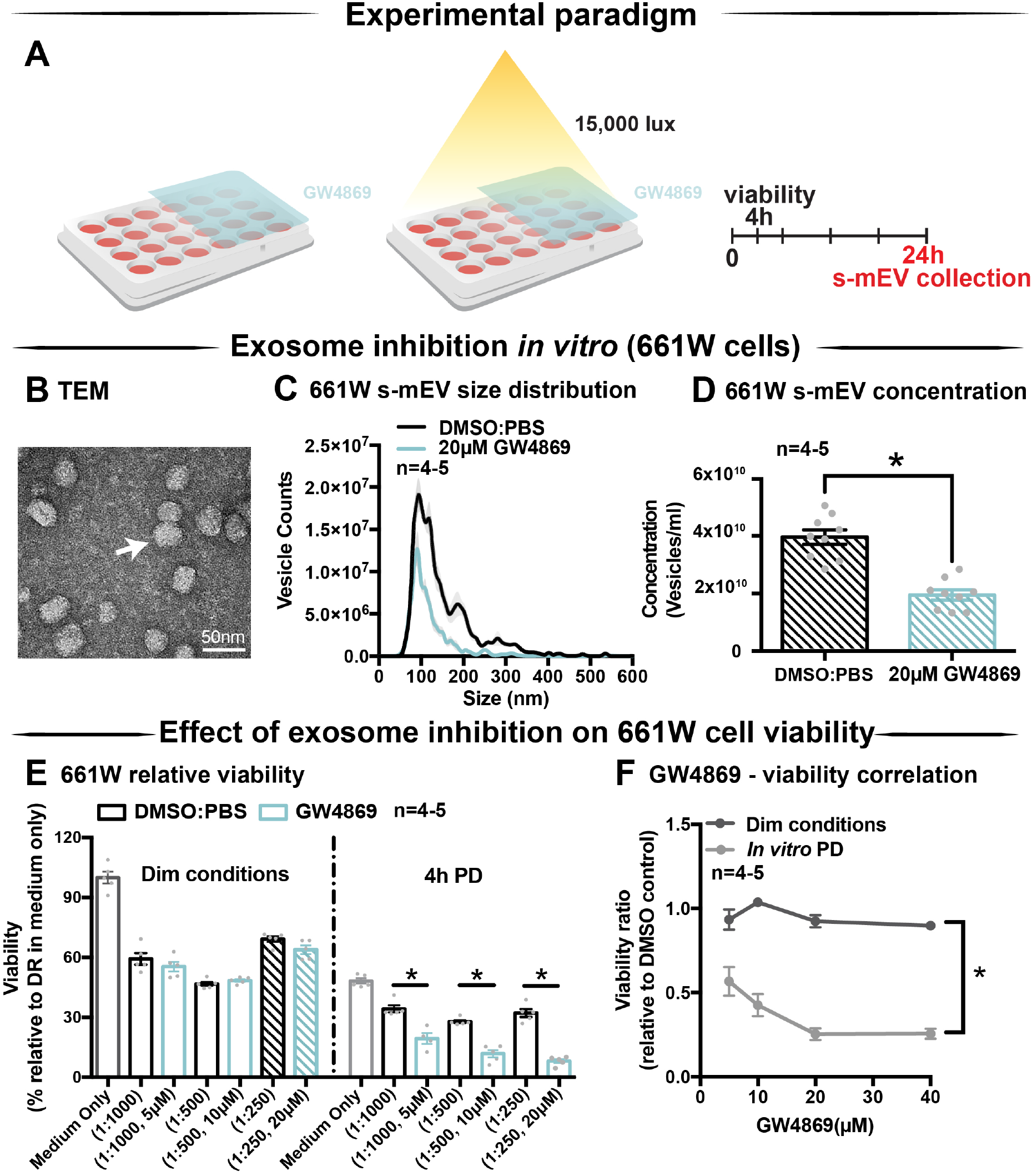
GW4869 inhibits bioavailability of s-mEV in vitro. **(A)** Experimental paradigm. **(B)** Representative TEM micrograph showing isolated 661W vesicles. **(C-D)** s-mEV isolated from 661W photoreceptor-like cells in the presence of exosome inhibitor GW4869 (20μM) or DMSO control for 24 hours showed a significant reduction in concentration, as measured by NanoSight NS300. (P<0.05, n=4-5). **(E)** MTT assay to determine viability of 661W cells treated with increasing concentrations of GW4869 or DMSO. Results demonstrated no dose response in viability levels for either treatment under dim conditions, however significant dose-dependent reduction in cell viability when treated with GW4869 under PD conditions (P<0.05). **(F)** The viability ratio (viability_GW4869_ / viability_DMSO_) of 661W cells treated with increasing concentrations of GW4869 in dim conditions or under 4 hours PD showed no change while in dim conditions, however is reduced between doses of 0-20μM GW4869 before reaching a plateau by 40μM while in PD conditions. Scale bar = 50nm. * Patterned bars in Figure 3E represent equivalent dosages/concentrations and conditions of GW4869/DMSO used in Figure 3D.

### 1.3.4. Endocytic pathway inhibition results in reduced s-mEV bioavailability *in vivo*

The contribution of exosomes to retinal s-mEV population was investigated *in vivo* (Figure Error*! No text of specified style in document.*.***4***A and B). s-mEV isolates from retinas of GW4869-treated mice (daily IP injections for 5 days) were reduced in concentration compared to those from DMSO-injected controls. This was seen in both dim-reared and photo-oxidative damaged retinas (Figure Error*! No text of specified style in document.*.***4***C and D). Additionally, the expression of genes associated with ESCRT-dependent and ESCRT-independent exosome biogenesis pathways were measured by qRT-PCR. We targeted *Pdcd6ip* (also known as *Alix*), which encodes an accessory protein in the ESCRT-dependent pathway, and *Smpd3*, which encodes nSMase2 in the ESCRT-independent pathway^207^. Retinal lysates from dim-reared and 5-day photo-oxidative damaged retinas as well as lysates from GW4869 and DMSO-injected photo-oxidative damaged retinas were used. The expression of *Pdcd6ip* and *Smpd3* was significantly increased in 5 day photo-oxidative damaged retinas compared to dim-reared controls (Figure Error*! No text of specified style in document.*.***4***E). Conversely, *Pdcd6ip* and *Smpd3* were found to be significantly reduced in GW486-injected retinas compared to DMSO-injected controls (Figure Error*! No text of specified style in document.*.***4***F). These results demonstrate that GW4869 can be used as an inhibitor of exosome production, and that exosomes likely contribute to the population as well as effects of retinal s-mEV.

**Figure Error! No text of specified style in document..4:**
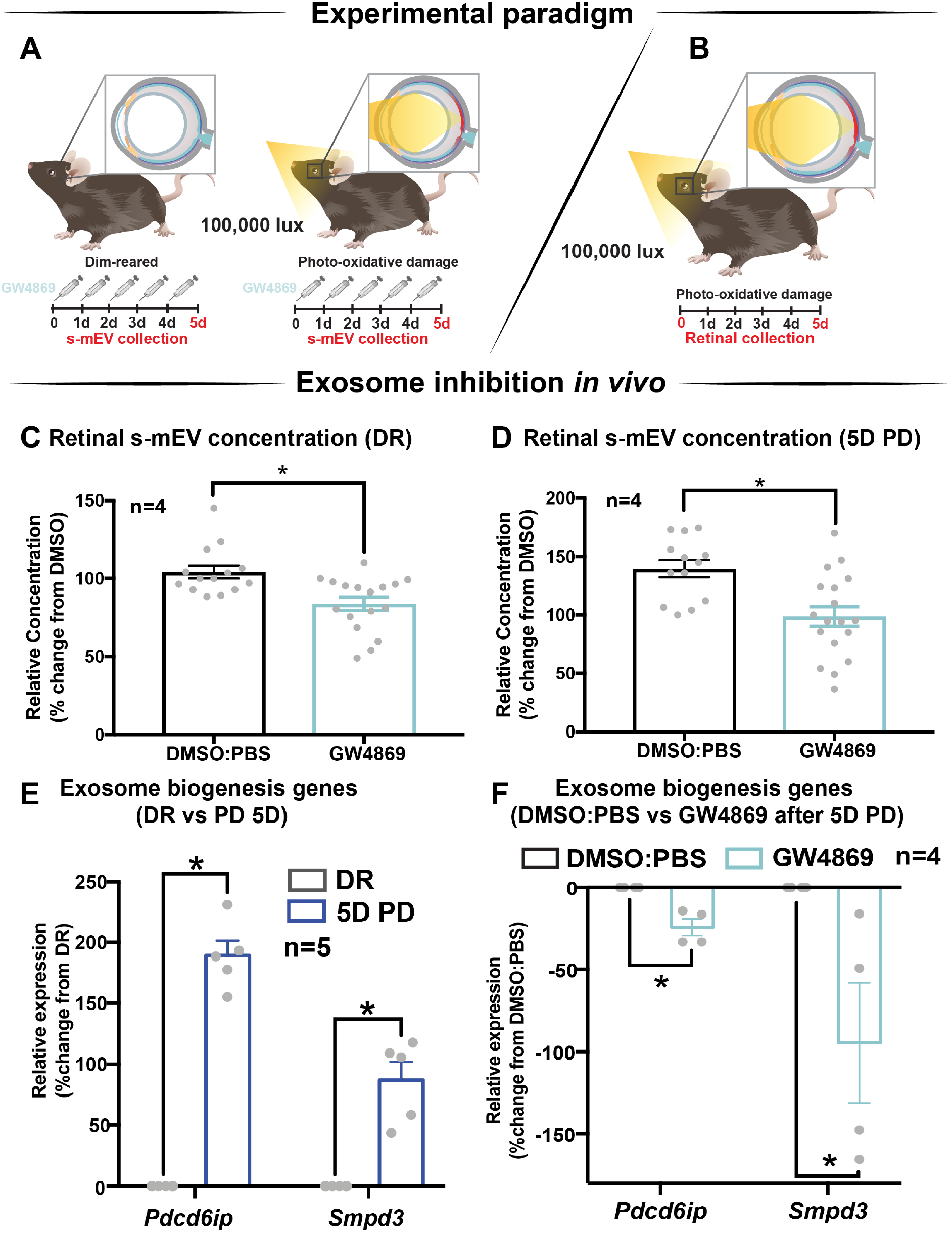
GW4869 inhibits bioavailability of retinal s-mEV. **(A-B)** Experimental paradigm with red retinal labelling indicating lesion site. **(C-D)** Nanotracking analysis (NanoSight NS300) quantification revealed a significant reduction in the concentration of s-mEV isolated from both DR and 5D PD retinas following 5 daily intraperitoneal injections of GW4869 compared to respective DMSO controls (P<0.05, n=4). The expression of exosome biogenesis genes *Pdcd6ip* and *Smpd3* as measured by qRT-PCR was **(E)** significantly increased following 5D PD relative to DR controls (P<0.05, n=5) and **(F)** significantly reduced following daily injections of GW4869 while undergoing 5D PD, relative to DMSO injected controls (P<0.05, n=4).

### 1.3.5. Exosome inhibition reduces retinal function in dim-reared mice

The effect of GW4869-mediated exosome inhibition on retinal health was investigated in dim-reared mice (Figure Error*! No text of specified style in document.*.***5***A). Retinal function was assessed using electroretinography (ERG). We show that GW4869-treated mice had reduced retinal function for both a-wave and b-wave measures (Figure Error*! No text of specified style in document.*.***5***B-D), compared to DMSO injected controls. To investigate this further, we measured photoreceptor cell death by counting TUNEL^+^ cells in the outer nuclear layer (ONL). The inflammatory response was also measured by counting IBA-1^+^ cells in the outer retina. No significant difference in either measure was observed between GW4869-injected and DMSO treated retinas (Figure Error*! No text of specified style in document.*.***5***E-G). Additionally, photoreceptor cell death was also assessed by measuring the ratio of ONL thickness to total retinal thickness from OCT images. No significant difference was detected between the two groups (Figure Error*! No text of specified style in document.*.***5***H-I). Overall, while GW4869 impaired retinal function in dim-reared mice, no difference was observed in histological measures between GW4869-injected mice and DMSO controls over the 5 day period investigated.

**Figure Error! No text of specified style in document..5:**
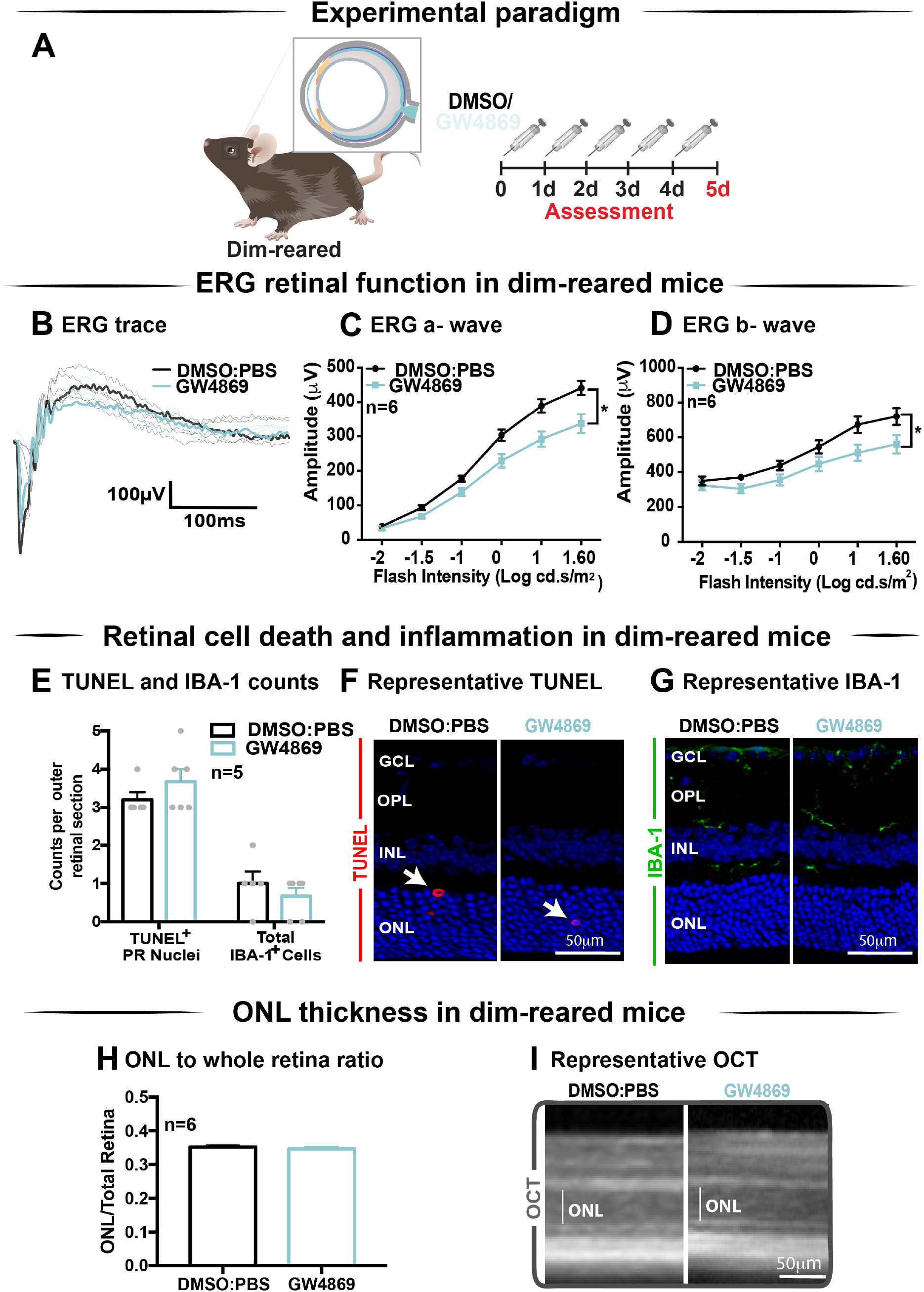
Exosome inhibition via the use of GW4869 in dim-reared mice reduces retinal function. **(A)** Experimental paradigm. Following 5 daily intraperitoneal injections of GW4869 (1.25mg/kg), retinal ERG function, **(B)** shown in representative ERG traces, was significantly reduced for both **(C)** a-wave and **(D)** b-wave responses, compared to DMSO controls in DR mice (P<0.05, n=6). **(E)** No significant changes were seen in either the number of TUNEL^+^ cells in the outer nuclear layer, or in the number of IBA-1^+^ cells in the outer retina following 5 daily intraperitoneal injections of GW4869 compared to DMSO controls (P>0.05, n=5). Representative confocal images show **(F)** TUNEL^+^ cells (white arrows) and **(G)** IBA-1^+^ cells in the outer retina, in both GW4869 and DMSO injected DR mice. Representative confocal images are taken in the superior outer retina, 1mm from optic nerve head. **(H)** There was no change in ONL thickness (represented as ratio to total retinal thickness), as measured by ocular coherence tomography (OCT) between GW4869 or DMSO injected DR mice (P>0.05, n=6). **(I)** Representative images of OCT scans show ONL thickness in superior retina, (white line) in GW4869 and DMSO injected DR mice. Scale bars = 50μM

### 1.3.6. Exosome inhibition reduces retinal function and increases cell death and immune cell recruitment in 5-day photo-oxidative damaged mice

Given the reduction in ERG responses in GW4869-treated dim-reared mice, the effect of exosome inhibition was further investigated in mice subjected to 5 days of photo-oxidative damage (Figure Error*! No text of specified style in document.*.***6***A). GW4869-injected 5 day photo-oxidative damaged mice showed significantly reduced retinal function for both a-wave and b-wave measures (Figure Error*! No text of specified style in document.*.***6***B-D), compared to DMSO controls. Interestingly, and contrasting to GW4869-treated dim-reared mice, we observed significantly higher levels of photoreceptor cell death (TUNEL^+^) and immune cell recruitment (IBA-1^+^) in GW4869-injected 5-day photo-oxidative damaged mice compared to DMSO controls (Figure Error*! No text of specified style in document.*.***6***E). The increased number of IBA-1^+^ cells in the outer retina included a higher ratio of amoeboid to ramified microglia and an increased presence of subretinal macrophages (Figure Error*! No text of specified style in document.*.***6***E). Further, there was a significant thinning of the ONL, indicating higher levels of photoreceptor cell death in GW4869-treated mice (Figure Error*! No text of specified style in document.*.***6***F). However, no difference in the thickness of the INL or GCL was observed (Figure Error*! No text of specified style in document.*.***6***F). Increased levels of photoreceptor cell death in GW4869-injected photo-oxidative damage mice were further demonstrated by a reduction of photoreceptor nuclei in the ONL (Figure Error*! No text of specified style in document.*.***6***G). Representative confocal images demonstrate an increase in photoreceptor cell death with significantly more TUNEL^+^ cells and thinner ONL (Figure Error*! No text of specified style in document.*.***6***H), as well as increased IBA-1^+^ cells (Figure Error*! No text of specified style in document.*.***6***I), in GW4869-treated mice compared to controls. Taken together these results show that exosome inhibition in retinas subjected to photo-oxidative damage causes reduced retinal function, concomitant with increased photoreceptor cell death and increased recruitment and activation of microglia/macrophages.

**Figure Error! No text of specified style in document..6:**
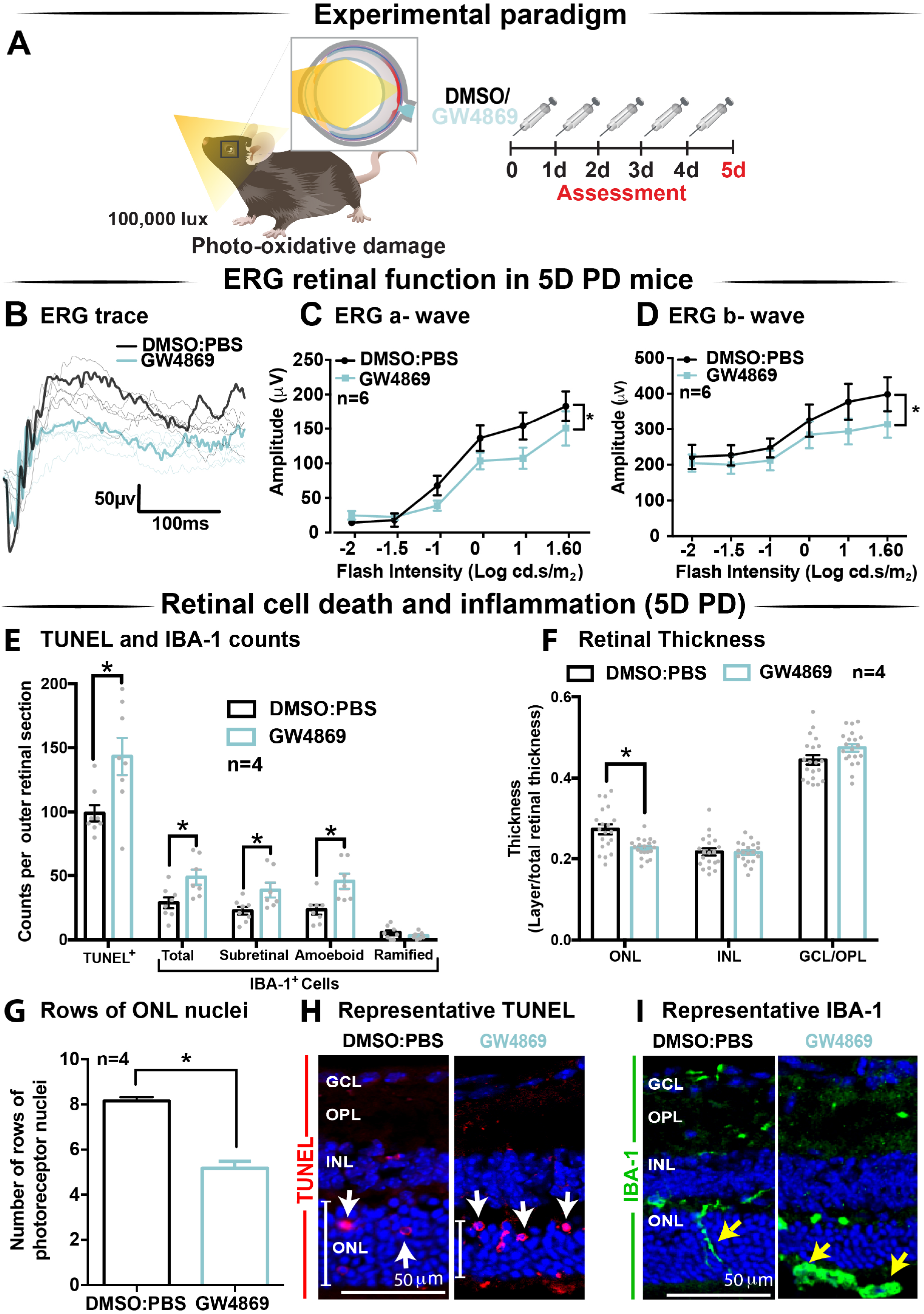
Exosome inhibition via the use of GW4869 in mice subjected to 5 days photo-oxidative damage reduces retinal function and increases cell death and immune cell recruitment. **(A)** Experimental paradigm with red retinal labelling indicating lesion site. Following 5 daily intraperitoneal injections of GW4869 (1.25mg/kg), retinal ERG function, **(B)** shown in representative ERG traces, was significantly reduced for both **(C)** a-wave and **(D)** b-wave responses, compared to DMSO controls in 5D PD mice (P<0.05, n=6). (**E)** There was a significant increase in the number of TUNEL^+^ cells in the ONL, as well as a significant increase in the total number, number of subretinal, and number of amoeboid but not ramified IBA-1^+^ cells in the outer retina in GW4869-injected mice compared to DMSO controls following 5D PD (P<0.05, n=4). **(F)** There was a significant reduction in the thickness of the ONL of GW4869-injected 5D PD mice compared to DMSO controls (P<0.05, n=4), however, no change was observed in the thickness of the INL or GCL between these groups (P>0.05, n=4). **(G)** The number of photoreceptor row nuclei in the ONL was also significantly reduced in GW4869-injected 5D PD mice compared to DMSO controls (P<0.05, n=4). Representative confocal images show **(H)** increased levels of TUNEL^+^ cells (while arrow) and reduced ONL thickness (white line) as well as **(I)** increased number of IBA-1^+^ cells in the outer retina, in GW4869-injected mice following 5 days of PD. Representative confocal images are taken in the superior outer retina, 1mm from optic nerve head. Scale bars = 50μM.

While small extracellular vesicle fractions isolated following high speed >100,000xg ultracentrifugation and expressing tetraspanin markers CD9, CD63 and CD81 (such as those isolated in this work) are commonly referred to as exosomes, without evidence of endosomal origin and in complying with MISEV 2018 guidelines ^238^, are herein referred to as small-to-medium EV, or s-mEV. In reference to other works, EV terminology will be referred to as published in the original papers.

### 1.3.7. Retinal s-mEVs possess a distinct miRNA profile compared to the retina which remains unchanged over 5-days of photo-oxidative damage

To obtain pure s-mEV-derived RNA, s-mEV isolated from dim-reared and 5 day photo-oxidative damaged retinas were treated with RNaseA prior to RNA isolation to remove potential contaminating RNA species attached to the outside. Bioanalyzer electropherograms show that RNaseA treatment successfully removed contaminating RNA, whereas s-mEV RNA was protected. Further, retinal s-mEV RNA was enriched for RNA species between 10 to 40 nucleotides (Supplementary Figure Error! No text of specified style in document.**.2**).

Databases of retinal and s-mEV miRNA were obtained using small RNAseq (Figure Error*! No text of specified style in document.*.***7***A), and were compared to determine if there was any selective enrichment of miRNA in isolated s-mEV. After generating normalized miRNA counts, a two-dimensional principal component analysis (PCA) was employed to visualize global differences between the expression of miRNA from dim-reared retinal s-mEVs and whole retinas. This analysis showed a clear segregation between the miRNA profiles of these groups (Figure Error*! No text of specified style in document.*.***7***B). Next, we examined the correlation between the expression levels of miRNAs from dim-reared retinal s-mEVs and whole retinas, finding a significant correlation (r=0.71, p<0.0001) between these groups. In particular, miR-183-5p, let-7b/c-5p, miR-124-3p, miR-125a/b-5p and miR-204-5p were highly expressed in both groups (Figure Error*! No text of specified style in document.*.***7***C). This correlation analysis also indicated that some miRNA species were enriched in s-mEVs over the retina, with differential expression analysis uncovering that 28 miRNAs are over-represented in s-mEVs (FC>+2, FDR<0.05), including miR-3770b, miR-706, miR-669c-3p, miR-7020-3p and miR-3770 which were all over 100-fold more abundant (Figure Error*! No text of specified style in document.*.***7***D).

**Figure Error! No text of specified style in document..7:**
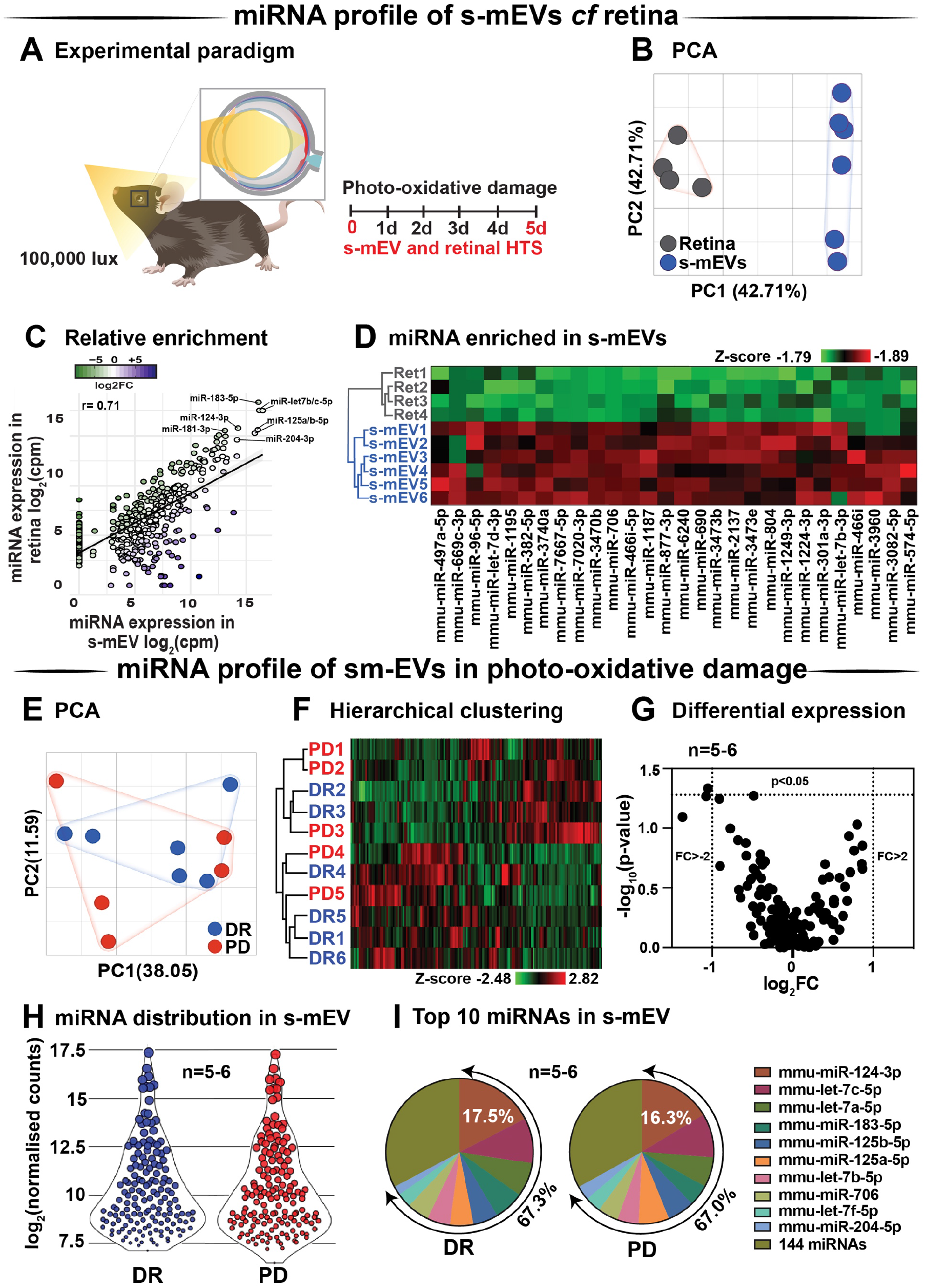
s-mEV miRNA profiling and enrichment analysis. **(A)** Experimental paradigm with red retinal labelling indicating lesion site. **(B)** Principal component analysis (PCA) of retinal miRNA and s-mEVs miRNA showed clear clustering suggesting that each group possesses uniquely enriched miRNA. **(C)** A positive correlation was observed (r = 0.71, p<0.0001) between miRNA counts in the dim-reared retina and s-mEVs with mir-124-3p, mir-181-3p, mir-204-3p, mir-125a/b-5p and miR-let7a/b-5p being abundant in both. **(D)** Hierarchical clustering of miRNA showed significant enrichment of miRNA in s-mEVs compared to the dim-reared retina (FC>2, FDR<0.05). **(E)** PCA and **(F)** hierarchical clustering analysis show no grouping or miRNA between DR and PD samples and identified no outliers. **(G)** Volcano plot indicating that no differential analysis was observed in s-mEV from DR and 5D PD retinas (P>0.05, n=5-6). **(H)** Violin plots showing the distribution of miRNA transcripts in s-mEV from DR and 5D PD retinas (n=5-6). Each data point represents the log_2_ transformed average number of counts for each miRNA retained after removing miRNAs with low expression and the size of each point is proportional to the average expression value of each miRNA. s-mEV from both DR and 5 days PD retinas contain similar amounts of miRNA transcripts with low, medium and high expression. **(I)** The top 10 most abundant miRNA were conserved in retinal s-mEV from DR and 5 days PD retinas and accounted for approximately 67% of the total counts.

Following, we examined the differential expression in miRNA between dim-reared and photo-oxidative damaged-isolated s-mEVs. PCA showed no clustering of the samples suggesting that the miRNA contents of retinal s-mEV were not substantially altered following photo-oxidative damage (Figure Error*! No text of specified style in document.*.***7***E). Hierarchical clustering analysis (HCA) further illustrated this, with no significant clustering between dim-reared and 5 day photo-oxidative damage groups (Figure Error*! No text of specified style in document.*.***7***F). In line with the PCA and HCA, differential expression analysis revealed that only miR-1249-3p was differentially expressed (p-value = 0.046, fold change = −2.08) (Figure Error*! No text of specified style in document.*.***7***G). However, when adjusting for multiple comparisons (Benjamini-Hochberg), this miRNA did not pass the significance threshold (Table S1).

A total of 154 miRNAs were detected in retinal s-mEV (Table S1) showing a similar mean count distributions in both groups (Figure Error*! No text of specified style in document.*.***7***H). The top 10 most highly expressed miRNAs were consistent in both dim-reared and 5 day photo-oxidative damaged retinal s-mEV, and accounted for approximately 67% of the total (Figure Error*! No text of specified style in document.*.***7***I). MiR-124-3p was the most abundant s-mEV-miRNA representing 17.5% and 16.3% in dim-reared and 5 day photo-oxidative damaged retinal s-mEV, respectively (Figure Error*! No text of specified style in document.*.***7***I). Four members of the let-7 family (let-7c-5p, let-7a-5p, let-7b-5p and let-7f-5p) as well as miR-183-5p, miR-125a-5p, miR-125b-5p, miR-706 and miR-204-5p comprised the rest of the top 10 most highly abundant s-mEV-miRNA (Figure Error*! No text of specified style in document.*.***7***I). These results demonstrate the selective enrichment of miRNA in s-mEV from the retina, but that the miRnome of s-mEV does not change in response to photo-oxidative damage.

### 1.3.8. s-mEV miRnome is associated with inflammatory, cell death and motility pathways

Considering that s-mEV miRNAs were not altered following 5 days of photo-oxidative damage and that the top 10 most abundant miRNAs accounted for 67% of the total s-mEV miRNAome, we focused on these top 10 s-mEV-miRNAs. A network analysis was performed to understand the interactions between the top 10 s-mEV-miRNAs and the retinal transcriptome (Figure Error*! No text of specified style in document.*.***8***A and Table S2). This was also performed for the enriched s-mEV miRNA. Network analyses of the top 10 most abundant s-mEV miRNA revealed that eight miRNAs form a regulatory network containing 1326 targets (Figure Error*! No text of specified style in document.*.***8***Bi), of which, miR-124-3p, mir-706 and let-7b-5p, have 472, 235 and 353 predicted interactions with retinal transcripts, respectively (Table S3). Similarly, the retinal targetome of the miRNA showing preferential enrichment in s-mEVs was explored with miRNet showing that miR-466i-3p, miR-446i-5p, miR-let-7b-5p, miR-1195 and miR-706 all had over 100 predicted targets in the retina (Figure Error*! No text of specified style in document.*.***8***Bii and Table S4). The predicted targets of both sets of miRNA were separately used for enrichment analyses against WikiPathways (mouse pathway annotation) and DisGeNET (database containing gene-disease associations) on the Enrichr platform. A total of 50 pathways were significantly over-represented in WikiPathways (Table S5 and S6). Notably, pathways pertaining to inflammatory processes including IL-1 to IL-6 signaling, Toll-like receptor signaling and chemokine signaling showed a significant enrichment. Pathways related to cell survival and motility were also significantly enriched (Figure Error*! No text of specified style in document.*.***8***C-D), with apoptosis significantly associated with miRNA targets from s-mEV-enriched miRNA (Figure Error*! No text of specified style in document.*.***8***C). Finally, DisGeNET database analysis revealed a significant association between s-mEV-miRNA targets and 6 retinal diseases (Figure Error*! No text of specified style in document.*.***8***E, and Table S7), with the majority associated with an inflammatory and/or proliferative profile. In summary, s-mEV miRNA cargo remains stable in the degenerating retina but the most abundant and enriched miRNA in s-mEV are predicted to modulate inflammation, cell death responses and cellular motility pathways.

**Figure Error! No text of specified style in document..8:**
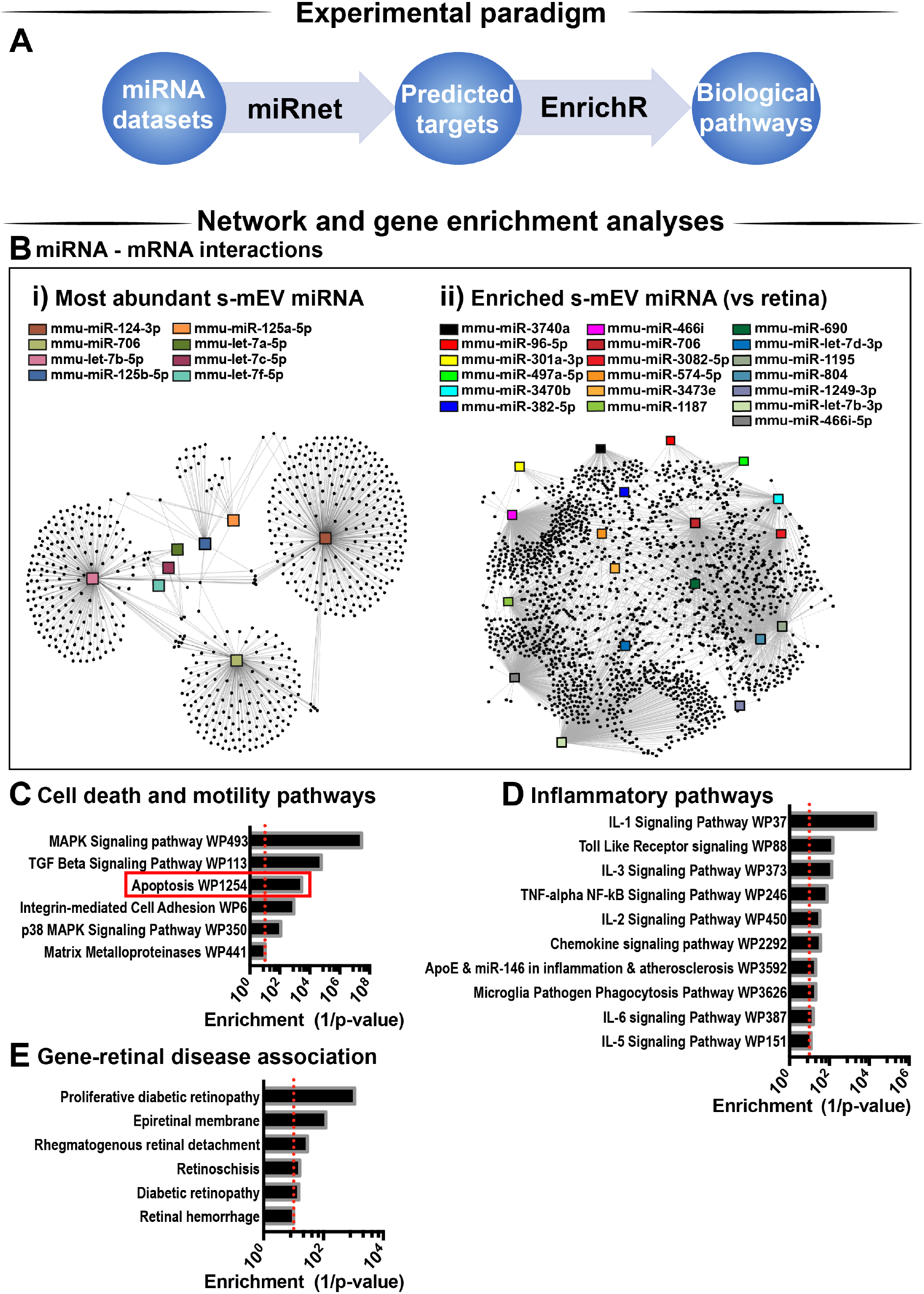
s-mEV miRNA target and pathway analysis. **(A)** Analysis pipeline. **(B)** miRNet network analysis using the **(i)** top 10 most highly expressed s-mEV-miRNAs and (ii) miRNAs enriched in s-mEVs showed their predicted interactions with target mRNAs. From the top 10 most abundant s-mEV miRNAs, eight formed an interconnected network, with miR-124-3p, miR-706 and let-7b-5p having the largest number of predicted targets. From the most enriched s-mEVs miRNAs, miR-466i-3p, miR-466i-5p and miR-1195 had the largest targetomes. Pathway analysis was performed using the predicted targets of the most abundant **(B, i)** and most enriched **(B, ii)** miRNA, collectively showing a significant enrichment of pathways pertaining to **(C)** cell survival and motility, **(D)** inflammatory processes and oxidative stress (P<0.05). **(E)** Enrichment analysis was also performed against DisGeNet (database containing gene-disease associations) revealing that a number of retinal diseases were significantly associated with the targets of s-mEV-miRNAs (P<0.05).

### 1.3.9. s-mEV may mediate the translocation of miR-124-3p from the outer to inner retina during photo-oxidative damage-induced retinal degeneration

We hypothesized that s-mEV may mediate the transportation of miRNA, including miR-124-3p, through the retina during damage. To test this, *in situ* hybridization of miR-124-3p was performed on retinal cryosections from mice injected daily with GW4869 and subjected to 5 days of photo-oxidative damage (Figure Error*! No text of specified style in document.*.***9***A). As previously reported^245^, miR-124-3p expression was significantly increased in the INL after 5 days photo-oxidative damage compared to dim-reared controls (Figure Error*! No text of specified style in document.*.***9***B-C). Interestingly, GW4869 treatment significantly reduced photo-oxidative damaged-induced miR-124-3p expression in the ONL and INL but not ILM/IS (Figure Error*! No text of specified style in document.*.***9***D-E). This indicates that the within-retinal expression, and potentially movement of miR-124-3p in response to photo-oxidative damage may be mediated by s-mEV such as exosomes. To further explore our hypothesis, we examined the expression of miR-124-3p in photo-oxidative damaged retinas after treatment with GW4869 and observed no differential expression compared to DMSO-injected controls (Figure Error*! No text of specified style in document.*.***9***F). This indicates that the reduced expression of miR-124-3p in the ONL and INL is likely due to lack of transport into the INL rather than a decrease in total retinal expression. Given the increased levels of cell death and inflammation in mice treated with GW4869, we speculate that s-mEV shuttling of miRNA including miR-124-3p is required for normal retinal homeostasis and immune modulation.

**Figure Error! No text of specified style in document..9:**
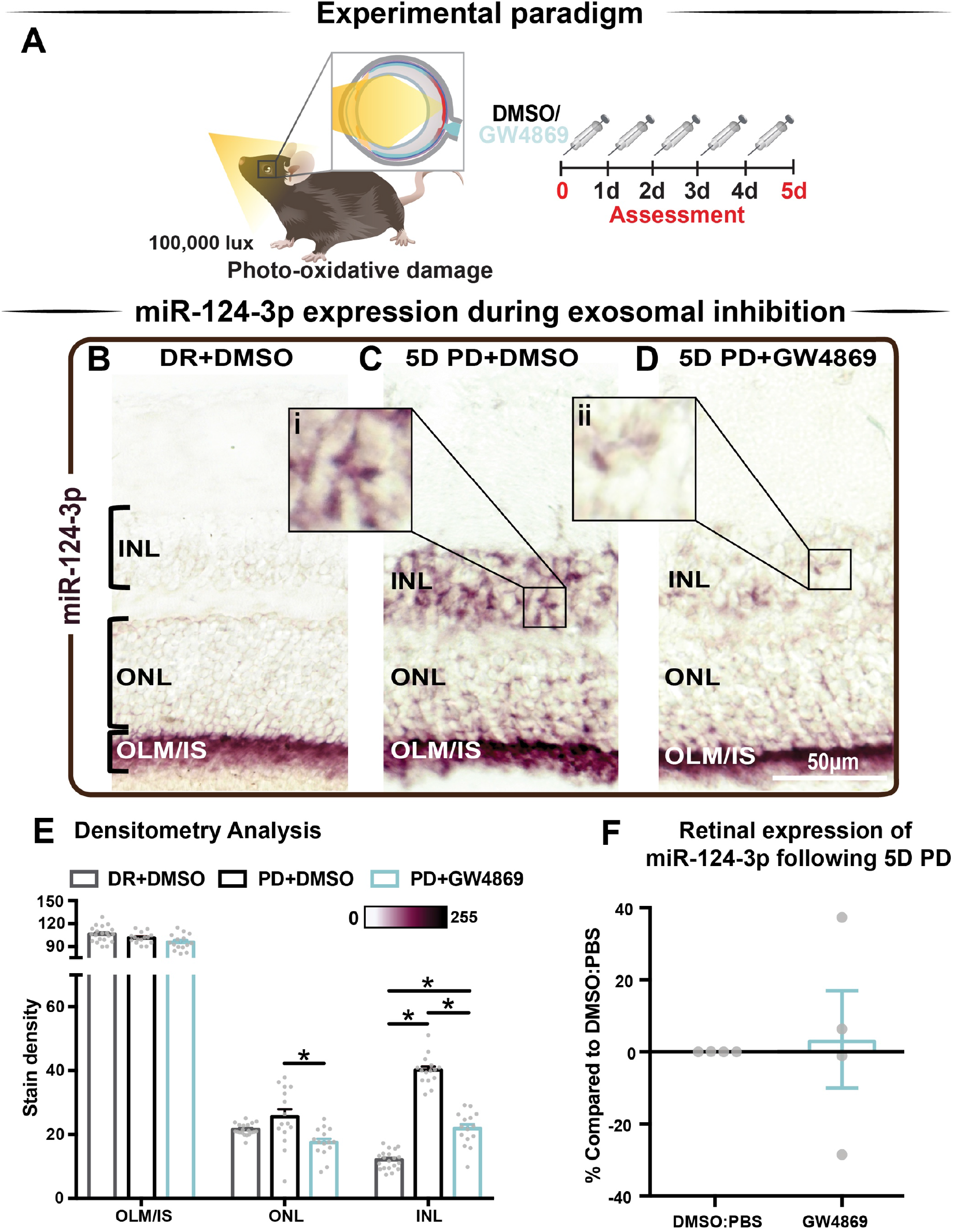
s-mEV may mediate the translocation of miR-124-3p from the outer to inner retina during photo-oxidative damage-induced retinal degeneration. **(A)** Experimental paradigm with red retinal labelling indicating lesion site. **(B)** Using *in situ* hybridization, miR-124-3p expression in DR retinal cryosections was localized to the outer limiting membrane/photoreceptor inner segments (OLM, IS), however **(C)** following 5D PD, miR-124-3p was seen in the INL (inlet, i). **(D)** Following intraperitoneal administration of GW4869 exosome inhibitor, miR-124-3p expression is significantly reduced in the ONL and INL (inlet, ii). Representative images are taken in the superior outer retina, 1mm from optic nerve head. **(E)** Densitometry analysis of miR-124-3p in retinal layers confirmed a decrease in miR-124-3p expression in the ONL and INL of mice treated with GW4869**. (F)** Retinal expression of miR-124-3p was conducted using qRT-PCR in photo-oxidative damaged mice treated with GW4869 or DMSO control, showing no change in total retinal expression (P>0.05). Scale bar = 50μM.

## 1.4. Discussion

This study describes for the first time, the isolation and characterization of mouse retinal s-mEV and the important role they play in retinal health and degeneration. We demonstrate four key findings from this study: Firstly, we demonstrate that s-mEV isolated from mouse retinas decrease in concentration progressively as a consequence of retinal degeneration. Secondly, we show that partial s-mEV depletion, via systemic administration of the exosome inhibitor GW4869, resulted in reduced retinal function in the normal retina and further exacerbated functional losses in mice subjected to photo-oxidative damage. Notably, significant photoreceptor cell death and inflammation were observed in GW4869-injected mice undergoing photo-oxidative damage, which was mirrored *in vitro* by 661W photoreceptor-like cells displaying increased susceptibility to photo-oxidative damage following exosome inhibition. Thirdly, we used small RNA sequencing and bioinformatic analyses to report on the potential regulatory roles of s-mEV miRNAs in retinal degeneration. Lastly, using the s-mEV-abundant miRNA miR-124-3p as a measure, we provide evidence that retinal s-mEV could be involved in the dynamic cell-to-cell transfer of miRNA in the degenerating retina. Taken together, we propose that retinal s-mEV and their miRNA cargo play an essential role in maintaining retinal homeostasis through immune-modulation, and that this effect is potentially mediated by s-mEV populations including exosomes.

### 1.4.1. Photoreceptor cell death is associated with reduced s-mEV bioavailability

Previous studies have demonstrated that miRNA-laden exosomes are found in abundance in the CNS and are central to cell-to-cell communication ^437,438^. Although it is unclear which retinal cell type(s) s-mEV isolated in this work are secreted by, our evidence supports a photoreceptor-derived contribution. We demonstrate a progressive decrease in s-mEV numbers during photo-oxidative damage, which correlates with increasing levels of photoreceptor cell death as previously described in this model ^265^. Therefore, we attribute the decrease in isolated retinal s-mEV to a loss of photoreceptors. In addition, the high abundance of photoreceptor prominent miRNAs miR-124-3p ^245^ and miR-183-5p ^439^ in isolated retinal s-mEV, further attests to a potential photoreceptor origin, with miR-124-3p comprising 17.5% and 16.3% of the total s-mEV-miRnome in normal and damaged retinas, respectively.

Previous studies by Vidal-Gil et al. (2019) report that the exosome protein marker CD9 was localized to the photoreceptor inner segments and ONL, further supporting a photoreceptor-origin hypothesis. However, it should be noted that CD9 was also found in the INL, GCL and choroid. Using a genetic model (*rd10* mice) of retinitis pigmentosa, which results in progressive rod photoreceptor-specific cell death ^440^, Vidal-Gil et al. (2019) showed that EVs isolated from retinal explants contain rhodopsin protein. This exosomal rhodopsin expression increased following photoreceptor rescue. Therefore, the authors describe EV secretion as a function of photoreceptor cell death, with significantly lower levels of CD9 expressing vesicles isolated from *rd10* mouse retinas at postnatal day 18 (P18) compared to at P16 ^415^. Taken together these results demonstrate a clear relationship between retinal s-mEV secretion and photoreceptor cell survival, suggesting a significant s-mEV population may derive from photoreceptors in the retina.

Contrary to our findings, it is well established that a pro-oxidative stress or a proinflammatory environment often results in the increased release of exosomes ^207,441-443^. We do observe increased expression of exosome biogenesis genes *Pdcd6ip* and *Smpd3* in retinal lysates following 5 days of photo-oxidative damage. We therefore suggest that all retinal cells including photoreceptors, as demonstrated by Vidal et al. (2019), may be potentially increasingly releasing s-mEV including exosomes as a consequence of the heightened presence of inflammatory and oxidative stress stimuli created in response to acute retinal degenerations. Further, we hypothesize that photoreceptors, as the most abundant retinal cell type, are the largest contributor to the pool of retinal s-mEV and thus their loss leads to a net decrease in the total bioavailability of retinal s-mEV. Independent isolation and quantification of s-mEV and in particular exosomes from each major retinal cell type may shed some light in this regard and will be investigated in the future. We also note that it is also possible that the increased expression of *Smpd3* may be occurring as a result of degeneration, with nSMase2/ceramide-induced apoptosis reported in models of retinal degenerations ^444-446^. While we do not see a protection from cell death following treatment with GW4869, we suspect that reduced exosomal communication may supersede possible protective effects from nSMase2/ceramide inhibition. Investigations into the interplay between exosome release and retinal ceramide levels will be explored in future works.

An additional hypothesis is that in response to excessive light, increased numbers of retinal s-mEV are mobilized as early responders to stress. However, even by 1 day of photo-oxidative damage, have already been depleted past a homeostatic level, ensuing an inflammatory response. As we previously demonstrated the potential translocation of miR-124-3p to the INL within 24 hours of damage ^245^, it seems possible that s-mEV transport of this miRNA could have occurred before this early time point as a rapid response to stress. In fact, ^447^ demonstrated that photoreceptor-specific miRNAs such as miR-183/182/96 as well as miR-204 and miR-211 are modulated in response to light exposure as short as 3 hours. We also identified mir-183/182/96, miR-204 and miR-211 in retinal s-mEV however, while Krol and colleagues do not propose a mechanism responsible for the miRNA turnover observed in their work, our work presented here suggests that s-mEV may be involved. Quantifying s-mEV secretion during early stress responses is required to determine if s-mEV depletion is a cause or consequence of retinal degeneration, and represents an important piece of the therapeutic puzzle.

### 1.4.2. Retinal s-mEV as mediators of immuno-modulation

Regardless of origin, to date the role that s-mEV play in retinal health and disease is still largely unclear. Results from this work strongly support a mechanism by which retinal s-mEV and in particular exosomes mediate homeostasis and immuno-modulation, with the inhibition of exosomes using GW4869 both *in vitro*, and *in vivo*, resulting in increased cell death, as well as recruitment and activation of microglia/macrophages. Importantly these observations were only evident under stress conditions, with both control 661W cells and dim-reared retinas displaying no major signs of cell death or inflammation following exosome-inhibition in the absence of photo-oxidative damage. We suggest that unlike in the degenerating retina, that exosome inhibition had no major effects on cell health. This is likely to be a consequence of the experimental paradigm used in this study and the short period of inhibition, or alternatively could indicate that under stress exosomal communication is necessary for cell survival. A smaller average size was seen in 661W-isolated s-mEV compared to those from the retina, and while we attribute this to the heterogenous nature of whole tissue, it could suggest that photoreceptors primarily secrete a smaller s-mEV fraction such as exosomes, and it is this population that may mediate retinal damage. This hypothesis however requires further investigation.

We hypothesize that as a consequence of longer-term exosome inhibition, inadequate translocation of miRNA cargo via s-mEVs results in the dysregulation of immune pathways. We have previously reported that in response to photo-oxidative damage, miR-124-3p upregulation in the INL may occur via outer-to-inner retinal translocation, with miR-124-3p acting as an anti-inflammatory regulator of C-C Motif Chemokine Ligand 2 (Ccl2) to prevent the recruitment of microglia/macrophages ^245^. In this present study, we provide further evidence of s-mEV-mediated miRNA translocation, demonstrating that in mice treated with GW4869, INL upregulation of miR-124-3p, the most highly expressed miRNA in isolated retinal s-mEV, was reduced. Although correlative, we suggest that insufficient gene regulation due to reduced s-mEV/miRNA bioavailability could contribute to the increased presence of immune cells and inflammation as seen in retinal degenerations. While we do not exclude the possibility that miR-124-3p could be upregulated in the INL in response to photo-oxidative damage, and downregulated following treatment with GW4869, we believe that this is unlikely, given the lack of differential change in the expression of miR-124-3p in photo-oxidative damaged mice injected with GW4869 compared to controls. The s-mEV-dependent transport of miRNA in mediating immune regulation however requires further exploration particularly in regard to the hypothesized movement of s-mEVs in retinal damage. Future experiments should utilize EV reporter strains such as those generated by Men et al, (2019) ^448^ as well as intra-ocular administration of fluorescently tagged s-mEVs to obtain direct evidence of s-mEV movement and uptake in the degenerating retina, and will be investigated in further detail in the future.

To further support the hypothesis that retinal s-mEV mediate homeostasis and immune responses, network and pathway analysis of retinal s-mEV-miRNAs revealed that the top 10 miRNA and miRNA enriched in s-mEVs were associated with the regulation of inflammatory and cell survival pathways. For example, our network analysis indicates that the top 10 miRNA are controlling genes associated with Interleukin and chemokine signaling, with both of these biological pathways involved in a plethora of retinal degenerative diseases ^53,62^. Moreover, MAPK signaling and TGF-β have also been shown to play pivotal roles in the development of retinal degenerations ^449,450^, with both pathways showing a strong association with the targetome of the top 10 most highly expressed s-mEV-miRNAs. As the top 10 miRNAs in both dim-reared and photo-oxidative damage retinal s-mEV make up approximately 70% of the total s-mEV-MiRnome, it is not surprising that a reduction in s-mEV numbers during degeneration could lead to immune dysfunction due to inadequate gene regulation. In fact, as recently demonstrated by Bian et al, (2020) exosomes secreted from transplanted neural stem/progenitor cells (NPCs) in the subretinal space resulted in delayed photoreceptor cell death, preserved retinal function and the suppressed activation of retinal microglia. Further, this group demonstrated that the miRNA cargo of NPC exosomes mediated this protection, with many NPC-derived miRNA found to be similar to s-mEV-derived miRNA reported in this study ^451^. Enrichment analysis of retinal s-mEV-MiR revealed significant associations with retinal degenerative diseases such diabetic retinopathy, retinal detachment and retinal hemorrhage further highlighting their potential involvement in modulating retinal inflammatory diseases.

From these collective findings, we propose that in normal retinal health, s-mEV are secreted from photoreceptor cells, and are released to the surrounding retina to maintain a homeostatic environment. However, following photoreceptor cell death, we hypothesize that immune responses are no longer able to be regulated due to reduced s-mEV numbers and the bioavailability of miRNA cargo; leading to the upregulation of inflammatory pathways, infiltration and activation of microglia/macrophages and progressive retinal cell death (Figure Error*! No text of specified style in document.*.*10*); characteristic features of retinal degenerative diseases ^11,62,452,453^.

**Figure Error! No text of specified style in document..10:**
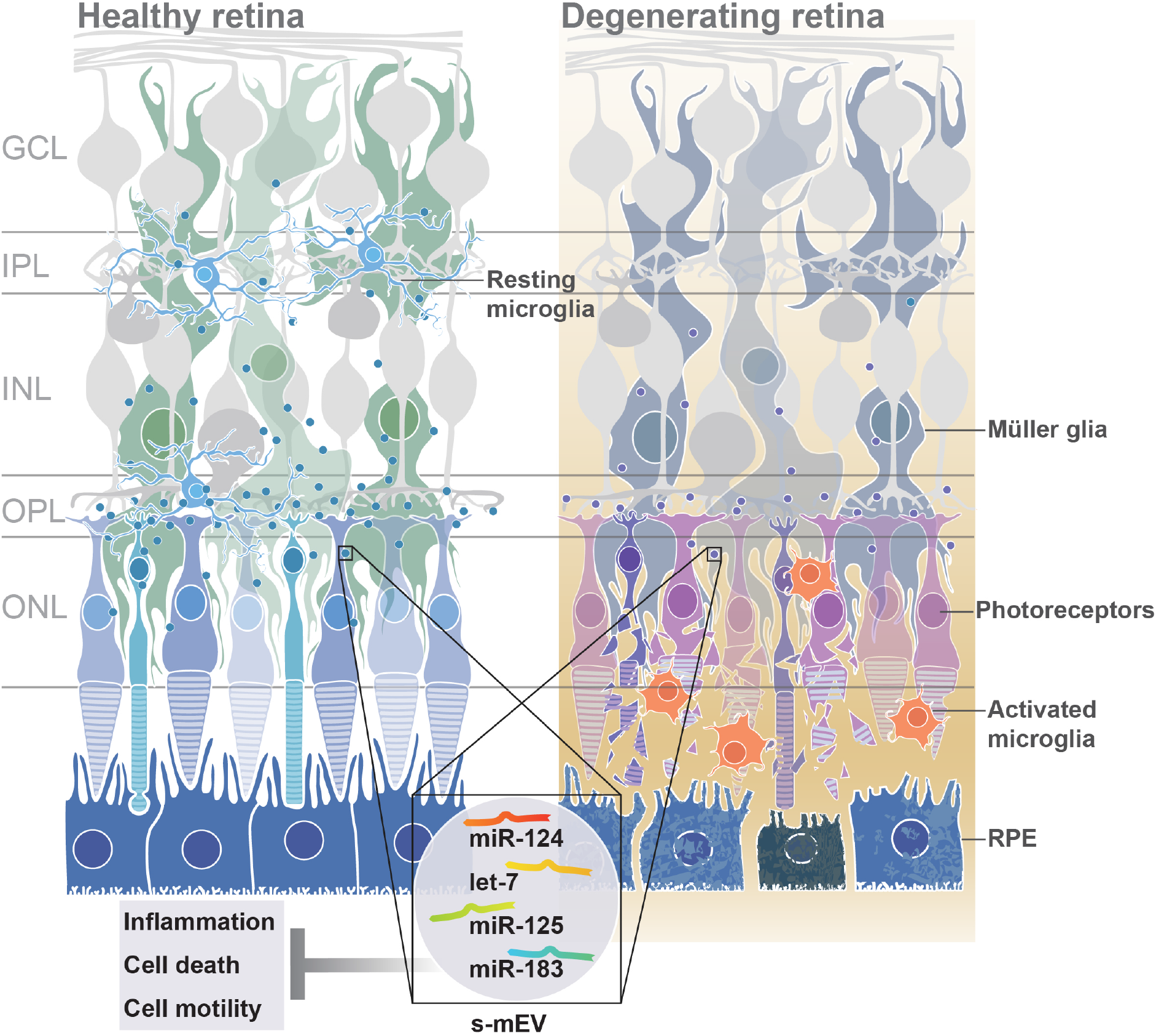
Proposed functional role of exosomes. Under normal homeostatic conditions, s-mEV laden with miRNA, including miR-124, let-7, miR-125 and miR-183, potentially originating from photoreceptors, are continuously trafficked between retinal neurons and glial cells. However, in retinal degenerations, a loss of photoreceptors leads to a decrease in s-mEV bioavailability, and consequently an insufficient transfer of s-mEV miRNA cargo to their cell targets. As the retinal targetome of s-mEV-miRNAs is associated with inflammation, survival and cell motility pathways, inadequate pathway regulation following s-mEV depletion leads to exacerbated retinal damage.

### 1.4.3. s-mEV as mediators of phototransduction

In addition to a potential role as regulators of retinal homeostasis through immune modulation, we highlight a possible involvement of retinal s-mEV, and in particular exosomes, in phototransduction. In both dim-reared and photo-oxidative damaged mice, exosome inhibition resulted in reduced retinal function as measured by ERG. While we could attribute the loss of retinal function in photo-oxidative damage mice to increased levels of photoreceptor cell death and immune cell recruitment, dim-reared mice showed no detectable signs of either cell death or inflammation at the time-point investigated, yet still demonstrated significantly lower functional responses.

While the role of s-mEV and exosomes in phototransduction in the retina is currently unknown, in the CNS the communication of exosomal cargo from oligodendrocytes and astrocytes has been reported to affect neurotransmission pathways^454-456^. In particular, Fruhbeis et. al (2013) suggested that the exosome communication between oligodendrocytes and neuronal axons is triggered by glutamate release ^457^. While oligodendrocytes are not present in the retina, Müller glia perform a similar role, responsible for the reuptake of glutamate released from photoreceptors terminals and the synaptic clefts of second order neurons as part of normal phototransduction ^458^. We speculate that, perturbations of exosomal communication systems could lead to aberrant glutamate reuptake by Muller cells and impaired signal transduction, explaining the observed reduction in retinal function of exosome-inhibited mice. However, this area requires further in-depth exploration.

### 1.4.4. The role of exosomes in retinal degenerative diseases and their potential as therapeutic gene therapy vehicles

To date, little progress has been made in uncovering the role that s-mEV may play in the progression of retinal degenerations, and further how we can utilize s-mEV, their cargo, and/or biological targets in diagnostic panels and in therapeutic development. It has been recently shown in cell culture models ^393,428,459^, that following high levels of oxidative stress (pathologically relevant to the onset of many retinal degenerative diseases), increased exosome numbers containing high expression of vascular endothelial growth factor receptor (VEGFR,) are secreted from a retinal pigmented epithelium (RPE) cell culture line (aRPE19) to promote neovascularization, a hallmark feature of several retinal pathologies ^428^. It has also been reported that exosomes containing retinal proteins were found in abundance in the vitreous humor of the mouse and human, demonstrating a level of communication that exists between these ocular tissues ^217^. While it has yet to be discovered if there is any differential change in exosome composition or concentration in the vitreous during retinal degenerative diseases, exosomes identified in the aqueous humor of AMD patients were found to contain a cross-over set of proteins also found in the culture medium of aRPE19 cells ^459^. This finding, while largely correlative, suggests that exosomes derived from the RPE may play a role in disease pathogenesis. Furthermore, these results demonstrate the possibility that access to exosome populations in biological fluids of the eye could provide a representation of what may be occurring in the retina and therefore serves a diagnostic potential.

Exosome based gene therapies are at the forefront of therapeutic development, with multiple clinical trials underway ^460,461^, including for the treatment of neurodegenerative diseases ^461,462^. However, their use for the treatment of retinal degenerations is largely in its infancy (reviewed in ^462^. Hajrasouliha et. al (2013) provides evidence for the immunomodulatory properties of exosomes derived from cultured retinal astrocytes ^463^. Using a laser induced model of choroidal neovascularization Hajrasouliha et. al (2013), demonstrated that the periocular injection of astrocytic exosomes reduced the CCL2-dependent migration of macrophages to the lesion site and attenuated angiogenesis ^463^. As CCL2 has been implicated as a key chemokine in the pathogenesis of multiple retinal degenerative diseases ^63,344,464^, and is a known target of miR-124-3p ^464^; exosome-based therapies that replenish retinal levels of this miRNA may prove efficacious as a possible therapeutic, as evidenced in other works ^245^. In addition, exosomes derived from microglial cells and injected into the vitreous of mice subjected to oxygen-induced retinopathy showed protective effects, reducing avascular regions in the retina, VEGF expression, and photoreceptor apoptosis, compared to controls ^465^. It was hypothesized by these authors that exosomal-miR-24-3p mediated this protection against hypoxia-induced cell death ^465^.

The combined findings of our work uncover a novel role for retinal s-mEV in both health and degeneration, unveiling a panel of s-mEV-miRNA required for retinal homeostasis, and target networks of these gene regulators comprising inflammatory, oxidative stress and cell survival pathways. As we elude to retinal health requiring optimal levels of s-mEV, and their cargo; replenishing s-mEV loads in the retina itself may prove as efficacious therapy, and will be the focus of future works. Further, both the unique s-mEV-miRNA signature and downstream target pathways open additional avenues for therapeutic development.

## 1.5. Conclusion

Results from this work suggest that s-mEV are released from photoreceptor cells to maintain retinal homeostasis. However, as a consequence of photoreceptor cell death, s-mEV secretion and/or bioavailability becomes reduced. Consequently, retinal s-mEV cargo, which contains regulatory miRNA and other molecules, are unable to regulate immune responses, subsequently contributing to progressive retinal cell death. We hypothesis that this mechanism is likely to be involved in many retinal degenerative and inflammatory diseases.

## 1.6. Supplementary Figures

**Supplementary Figure Error! No text of specified style in document..1:**
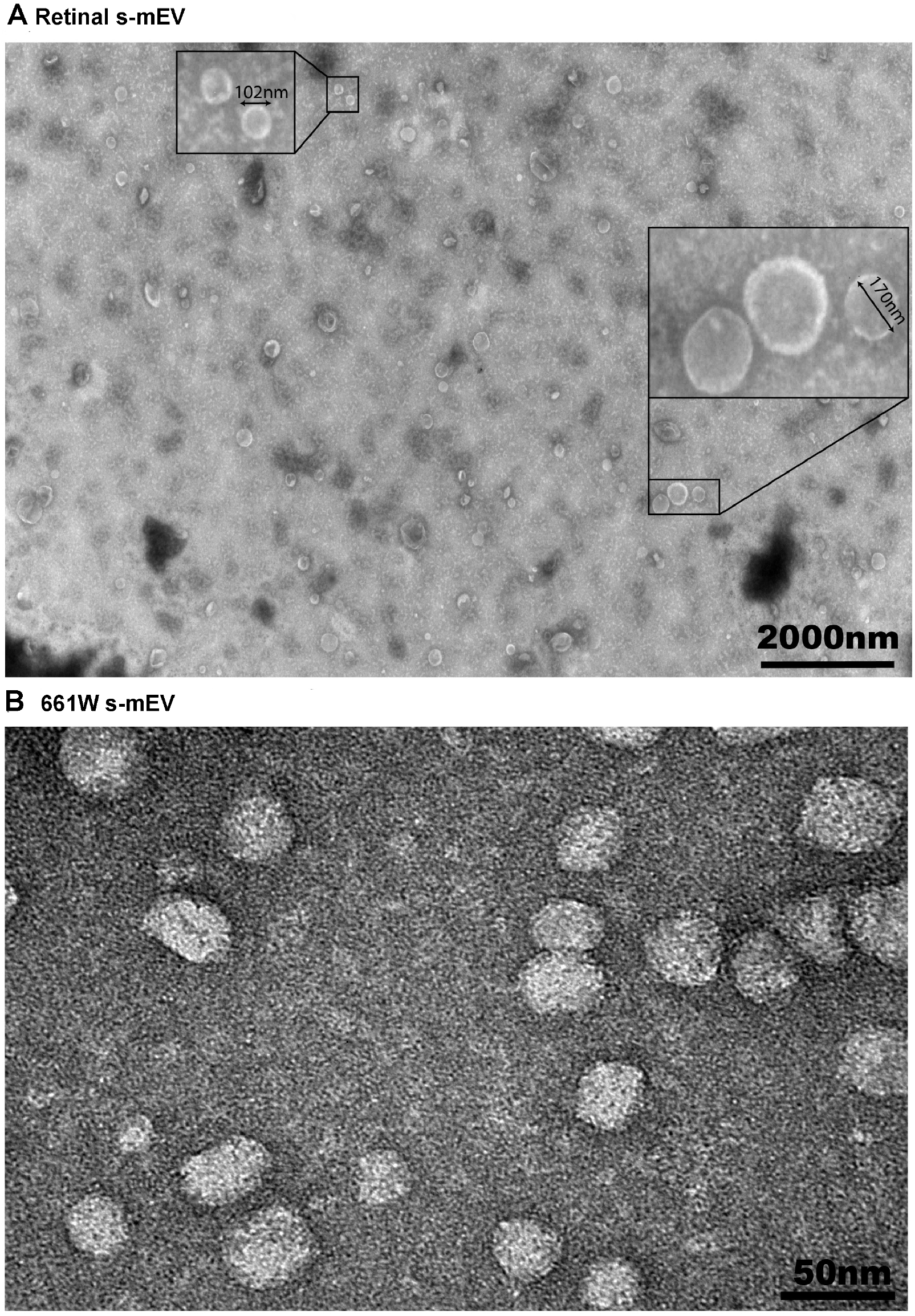
TEM. Representative TEM image showing isolated extracellular vesicles from **(A)** retinas and **(B)** 661W cells. Enlarged fields highlight the circular, cup-shape morphology and size of s-mEV such as exosomes. Scale bar = 2000nm and 50nm.

**Supplementary Figure Error! No text of specified style in document..2:**
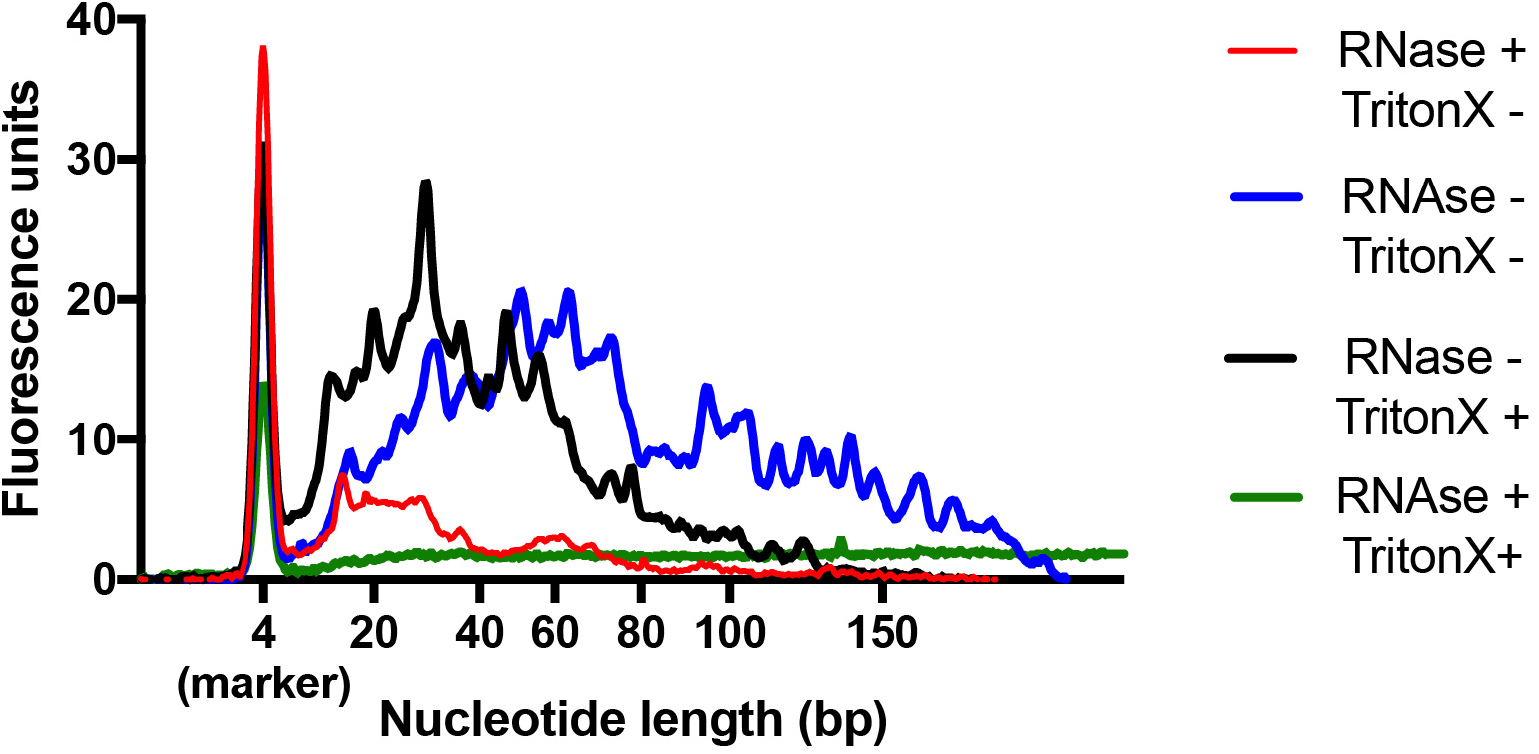
RNaseA Treatment. s-mEV were treated with RNaseA with or without Triton-X 100 for 30 mins then processed immediately for RNA extraction. Exosomes treated with RNAaseA without Triton-X 100 (red line) showed a decrease in the total amount of RNA compared with untreated s-mEV (blue line), and total RNA (black line), indicating the presence of contaminating RNA. The RNA protected from RNaseA degradation (red line) was enriched for RNA species of 10 to 40 nucleotides (bp) in length. The simultaneous treatment with RNasA and Triton-X 100 (green line) completely ablated the signal, further demonstrating that s-mEV-RNA is protected from RNAseA degradation.

**Supplementary Figure Error! No text of specified style in document..3:**
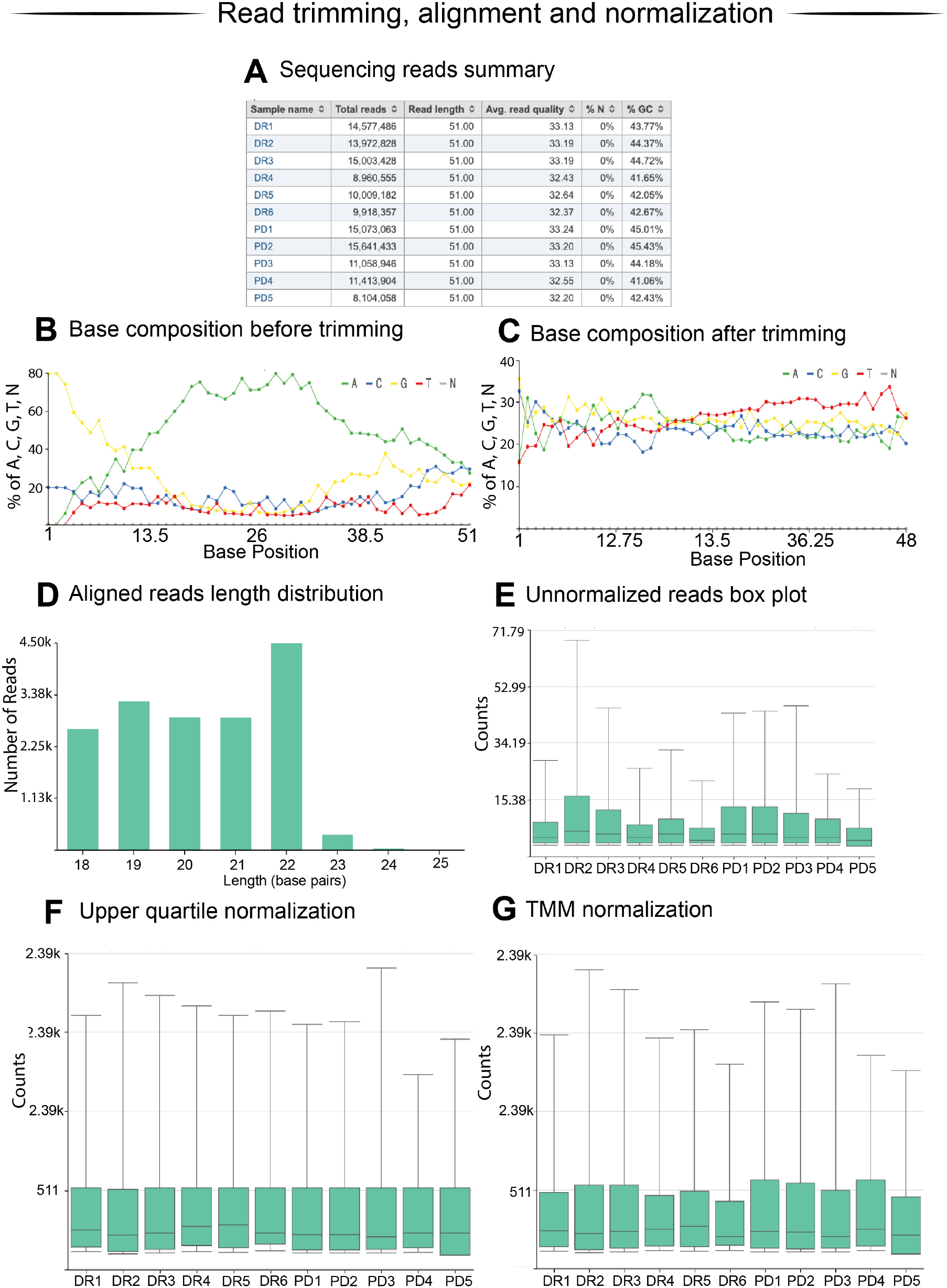
Sequencing alignment. **(A)** Summary of the sequencing depth, read quality and GC content**. (B)** Representative raw read base composition shows the presence of an enriched poly-A tail (green line) and template switching nucleotide (first 3 base pairs). These features are incorporated during the CATS library preparation and thus their enrichment in the sequenced reads indicates the successful construction of the library. **(C)** After trimming, every nucleotide was equally represented along the sequenced reads indicating the successful removal of library contaminants. **(D)** Reads aligning to miRbase v.22 (mature miRNA) had length distribution ranging from 18 to 24 nucleotides consistent with the length of annotated miRNAs. **(E)** After annotation, unnormalized counts were normalized using the **(F)** upper quartile (UQ) normalization or **(G)** Trimmed Means of M, with the former producing more similar means and distributions (each box-and whisker plot corresponds to a sample).

